# CD388: A universally protective Drug-Fc Conjugate that targets influenza virus neuraminidase

**DOI:** 10.1101/2024.06.04.597465

**Authors:** Simon Döhrmann, James Levin, Jason N. Cole, Allen Borchardt, Karin Amundson, Amanda Almaguer, Elizabeth Abelovski, Rajvir Grewal, Douglas Zuill, Nicholas Dedeic, Grayson Hough, Joanne Fortier, Joanna Donatelli, Thanh Lam, Zhi-Yong Chen, Wanlong Jiang, Travis Haussener, Alain Noncovich, James M. Balkovec, Daniel C. Bensen, Voon Ong, Thomas P. Brady, Jeffrey Locke, Jeffrey L. Stein, Leslie W. Tari

## Abstract

The ability of the influenza virus to elude humoral immunity by rapid antigenic shift presents a sustained and urgent threat to human health. Globally, seasonal influenza causes an estimated 3 - 5 million cases of severe disease and 300,000 - 500,000 deaths annually, with increased potential for mortality during pandemics^1^. The recent outbreaks of avian H5N1 in bird populations, subsequent spread to cattle and appearance in humans, highlight the need for effective broad-spectrum influenza antivirals for treatment, prophylaxis and pandemic preparedness. The narrow, strain-specific immunity induced by current seasonal vaccines and mismatches between vaccine strains and circulating viruses result in limited vaccine effectiveness (VE) rates^2^. Furthermore, VE is even lower in immune-compromised and - senescent populations^3^. Strategies that provide durable, universal influenza protection in healthy and high-risk populations are urgently needed. Here, we describe CD388, a first-in-class antiviral drug-Fc conjugate (DFC) in clinical trials for seasonal influenza prevention (NCT05285137 and NCT05523089). CD388 comprises a multivalent small molecule inhibitor of influenza virus neuraminidase (NA) linked to a C_H_1-Fc hybrid domain of human IgG1 engineered for extended half-life. CD388 demonstrated potent, universal activity across influenza A and B viruses, including high pathogenicity and NA resistant strains, a low potential for resistance development, and efficacy in lethal mouse infection models. CD388 is the first therapeutic with the potential for universal prevention of influenza A and B in healthy and high-risk populations.

## INTRODUCTION

Two major surface glycoproteins on influenza are targets for current vaccines and antiviral therapies – hemagglutinin (HA), a sialic acid binding glycoprotein essential for viral entry and fusion, and neuraminidase (NA), an enzyme that cleaves the terminal sialic acid from *N*-linked glycans, mediating viral egress^1,4^. Current vaccines primarily prevent influenza infection via induction of antibodies targeting the immunodominant antigenic sites on the HA head domain to block binding of influenza to its sialic acid receptor^5^. However, pressure from the immune system and error-prone genome replication result in a high mutation rate in the virus^6^. Accordingly, vaccines are generally strain-specific, necessitating annual vaccination with multivalent vaccines.

Antibodies raised against the more conserved but less immunogenic HA stalk domain demonstrate broad coverage across influenza A that encompasses group 1 and, in some cases, group 2 viruses. However, this coverage does not extend to influenza B^7,8^. To achieve greater breadth of coverage for both prevention and treatment of influenza, NA has long been recognized as an attractive target^9,10^. The NA enzymatic activity plays two key roles in the establishment of influenza infection and proliferation: the release of nascent virus particles budding from infected cells and the clearance of decoy sialic acids on mucin lining the respiratory tract that trap incoming virus particles before they reach their targets^9^. Structurally and compositionally, the active sites of NA are highly conserved across influenza A group 1 and 2 viruses and both influenza B lineages^11^. Additionally, genetic drift in NA is generally slower and discordant with that of HA^12,13^. These favourable attributes spurred efforts to discover effective, NA-targeting vaccines and monoclonal antibodies (mAbs) ideally with greater breadth of coverage than therapies targeting HA^9,14^. Indeed, the high degree of active site conservation, coupled with the unique active site architecture of influenza NA, even compared to intracellular human sialidases^15^, has been successfully leveraged to develop selective small molecule influenza NA inhibitors (NAIs) with broad-spectrum viral coverage that have become mainstays of influenza treatment for the past 20 years^16^.

Globally, three NAIs are licensed for treatment of influenza: zanamivir (ZAN, Relenza®) was the first-in-class^17^, followed by oseltamivir (OST, Tamiflu®)^18^, and then peramivir (Rapivab®)^19^. ZAN was designed based on the NA transition state analogue 2,3-dehydro-2-deoxy-N-acetylneuraminic acid decorated with a guanidine group at the C4 position to fill an empty negatively charged pocket in the active site. ZAN demonstrated best-in-class, balanced potency against influenza A and B neuraminidases compared with the other NAIs that were less potent against influenza B. However, the polar character of ZAN results in poor oral absorption and rapid renal clearance following intravenous (IV) administration, necessitating dosing by buccal inhalation, the consequence of which is increased risk of bronchospasms in patients on ventilators or with asthma or chronic obstructive pulmonary disease^20^. As a result, clinical use of ZAN has been limited despite its clear advantages over other approved NAIs. Notably, a high-dose, twice-daily, IV formulation of ZAN to treat patients with severe influenza on a compassionate-use basis was recently approved^21^. OST was designed based on knowledge garnered from ZAN and incorporation of several changes to improve oral absorption and drug-like properties, including removal of the C4 guanidine group and replacement of the glycerol substituent at C6 in ZAN with a bulky, lipophilic pentyl ether side-chain^18^. As the only oral NAI, OST has dominated in the treatment of uncomplicated influenza; however, OST is also the least potent of the three approved NAIs, particularly versus influenza B strains, and the most susceptible NAI to escape mutations^16^. The very modifications that improved the OST drug-like properties relative to ZAN altered the mechanism by which it engages the NA active site, inducing a conformational change to accommodate the bulky lipophilic pentyl ether^22^. Viral mutants that impede this conformational change confer high level resistance to OST and have emerged in the clinic with relatively high frequency^16,23^. ZAN by contrast, resembles the transition state conformation of the NA substrate, and does not require a conformational change for binding. Accordingly, OST-resistant influenza mutants show low cross-resistance to ZAN, and moreover, the incidence of ZAN-resistant mutants in the clinic is exceedingly rare^16^. Recently, baloxavir marboxil (BXM, Xofluza®), a cap-dependent endonuclease inhibitor, was approved for influenza treatment. BXM exhibits modestly improved clinical efficacy compared to OST. However, during clinical trials, treatment-emergent variants with reduced susceptibility to BXM were identified^24^.

Based on a comprehensive analysis of influenza targeting small molecule inhibitors and viral target conservation, ZAN was identified as an optimal starting point for the development of new antiviral agents, despite the weaknesses that limit its clinical use. To leverage and enhance the best-in-class activity of ZAN and simultaneously improve its pharmacological attributes, we developed a new therapeutic modality, a drug-Fc conjugate (DFC), represented by CD388, comprised of a stable, multivalent conjugate of a novel ZAN dimer with an N-terminal extended Fc domain of human IgG1 **(Fig. 1a)**, engineered to extend half-life. The multivalent presentation of ZAN dimers on CD388 conferred potent, universal activity in NA enzyme inhibition studies and cell-based assays against a panel of over 50 influenza A and B viruses. Importantly, CD388 demonstrated multi-log increased potency compared to OST or ZAN in cytopathic effect (CPE) assays and, crucially, CD388 retained activity against OST- and ZAN-resistant influenza strains. CD388 also conferred a high barrier to resistance development in serial passage studies conducted using sub-inhibitory and escalating CD388 concentrations. Moreover, the engineered Fc domain in CD388 demonstrated a long circulating half-life, and the smaller size of the conjugate (compared with a full-length mAb) resulted in rapid and robust distribution of CD388 to the lung and epithelial lining fluid (ELF) in mice. These features delivered potent efficacy in numerous lethal challenge treatment and prophylaxis mouse models. Single intramuscular (IM) doses of CD388 at 1 mg/kg or less resulted in full protection of mice against numerous influenza A and B strains. Notably, CD388 demonstrated dose-dependent reductions in viral lung burden and inflammatory cytokines. CD388, currently in clinical development for prevention of influenza (NCT05285137 and NCT05523089), is a new therapeutic modality with the potential to address the longstanding unmet need for durable, universal treatment and long-term prevention of both seasonal influenza and influenza strains with pandemic potential.

**Figure 1.**
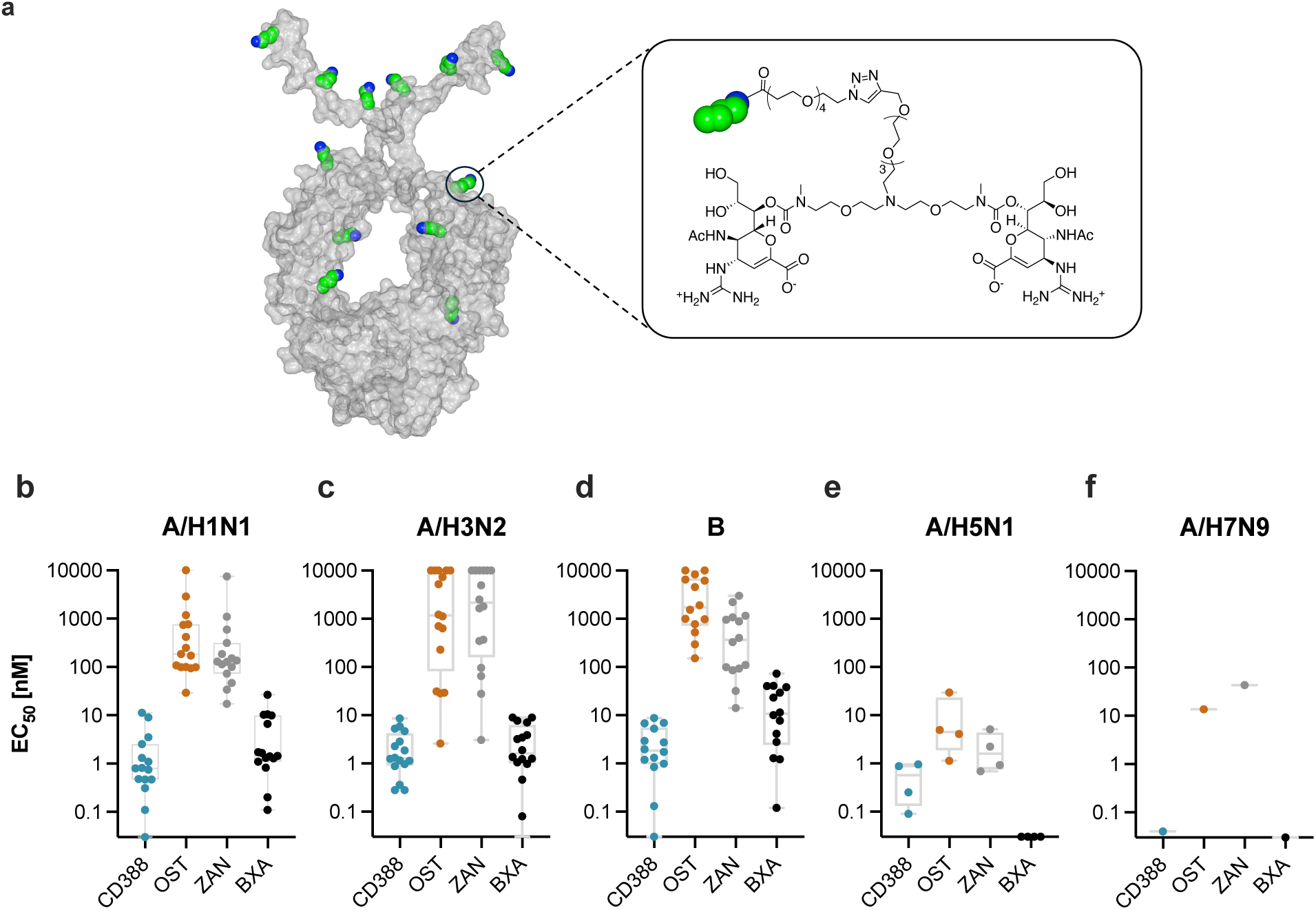
Structure of CD388 and universal activity against influenza A and B in cell-based CPE and microneutralization assays. (**a**) Molecular surface representation of the full-length hIgG1 (PDB code 1HZH) structure redacted to show only the Fc construct boundaries used in CD388. Positions of lysine residues that were preferentially conjugated in CD388 are shown in green (based on proteolytic digestion and peptide mapping using a conjugate with wild-type hIgG1; **Supplementary methods**). CD388 is a multivalent conjugate of a ZAN dimer (shown in blowup) stably conjugated to lysines on the N-terminal extended hIgG1 Fc with an average drug-antibody ratio (DAR) of 4.5:1. The spatial disposition of conjugated lysine residues on the Fc domain surface and the length and flexibility of the PEG linkers result in ZAN dimer constellations on CD388 where sufficient separation (> 90 Å) exists between ZAN dimers to allow simultaneous engagement of a single CD388 molecule with multiple active sites within an NA tetramer, between neighboring NA tetramers on the same virus particle, or between NA tetramers on different virus particles. EC_50_ values [nM] of CD388, OST, ZAN or BXA against a panel of influenza strains (**b**) A/H1N1, (**c**) A/H3N2, (**d**) B in a cell-based cytopathic effect assay, and (**e**) A/H5N1 and (**f**) A/H7N9 in a cell-based microneutralization assay.

## RESULTS

### Concept and design of CD388

Multivalent interactions (i.e., interactions between multiple ligands on one molecule and multiple receptors residing on a second moiety) are employed frequently in biological systems. They can serve to aggregate receptors to amplify signalling and dramatically increase the avidity of important functional interactions^25^. The homotetrameric structure of NA (**Extended Data Fig. 1**) and the dense clustering of NA on multiple regions on the surface of the influenza viral envelope^26,27^ make influenza an ideal target for multivalent NAIs. Previous research has shown that dimeric ZAN compositions using linkers that space the ZAN monomers (optimally) by 18 to 22 Å demonstrate improved antiviral activity when compared to monomeric ZAN in cell-based assays and mouse infection models^28^. While the ZAN spacing in dimers was insufficient to bridge adjacent monomers in the NA tetramer, the ability to promote aggregation of virus via “head-to-head” cross-linking of isolated NA tetramers on different virions was correlated with enhanced antiviral activity^28,29^. More recently, tetrameric ZAN compositions with larger spacing between the ZAN moieties, sufficient to bridge all four NA monomers within a NA tetramer, demonstrated marked improvements versus monomeric ZAN in binding affinity to NA, and drove improvements in in vitro and in vivo antiviral activity^30^. However, in both examples above, the inhibitor candidates did not overcome the IV administration route limitations of ZAN to justify advancement to the clinic.

CD388 contains chemically and metabolically stable ZAN dimers conjugated to an N-terminally extended hIgG1 Fc domain. The Fc domain has two modifications; a C220S mutation near the N-terminus to reduce the propensity for disulfide-bond-mediated aggregation, and the M252Y/S254T/T256E (YTE) triple mutation at the C_H_2-C_H_3 interface which increases half-life^31^. The ZAN dimers are symmetrically fused through N-methyl carbamate moieties on the C7-hydroxyl of ZAN to a flexible 15 atom central-linker that separates the ZAN monomers by approximately 18 Å. The carbamate linkage to ZAN was N-methylated to prevent spontaneous rearrangement of the carbamate from the secondary hydroxyl on C7 to the primary hydroxyl at the C9 position. Multiple ZAN dimers are heterogeneously conjugated to surface-exposed lysine residues on the Fc domain through a 32-atom flexible polyethylene glycol (PEG)-based linker, which projects them out from the Fc surface by 35 - 40 Å when the linker is fully extended (**Fig. 1a**). Specific solvent-exposed lysine residues were preferred sites for conjugation and the average ratio of ZAN dimer drugs to Fc in CD388 of 4.5:1 provides antiviral activity and favorable solution properties. Synthesis of the ZAN dimer and final conjugate are described in the **Supplementary Methods**. The spatial distribution of preferred sites for conjugation coupled with the length and flexibility of the cross-linker allows CD388 to simultaneously interact with multiple NA active sites within a tetramer (ca. 45 Å or 70 Å) (**Extended Data Fig. 1**). This arrangement has previously been shown to improve potency for tetrameric ZAN^30^. Additionally, the spacing of ZAN dimers allows bridging of NA active sites across neighbouring NA tetramers on the same virion (25 - 30 Å) or across two virions (16 Å)^28^. Thus, CD388 has the potential for multivalent engagement of NA combined with the potential to sterically interfere with NA-host cell interactions and promote viral aggregation.

### Universal activity of CD388 against influenza

We investigated the intrinsic target-based activity of CD388 to inhibit viral NA in a NA inhibition assay. CD388 demonstrated potent activity against all influenza A and B strains tested. CD388 had a median IC_50_ of 1.29 nM (*n* = 17; ranging from 0.01 to 2.36 nM) against subtype A/H1N1, 2.24 nM (*n* = 18; ranging from 0.31 to 3.88 nM) against subtype A/H3N2, and 2.37 nM (*n* = 13; ranging from 0.05 to 7.44 nM) against influenza B (**Table 1**). In general, CD388 performed similarly to ZAN and OST but was more potent than OST against influenza B strains (**Table 1**). Moreover, CD388 demonstrated potent activity against recombinant NA derived from highly pathogenic avian influenza (HPAI) viruses A/H5N1 and A/H7N9 with an IC_50_ of 3.72 nM (*n* = 1) or a median IC_50_ of 1.23 (*n* = 2; ranging from 1.03 to 1.43 nM), respectively (**Table 2**). OST and ZAN had comparable activity versus recombinant NA from A/H5N1 and A/Anhui/1/2013 (H7N9), but OST and ZAN lost substantial activity against NA from A/Shanghai/1/2013 (H7N9) with IC_50_ values of >1 µM and >100 nM, respectively (**Table 2**).

**Table 1.**
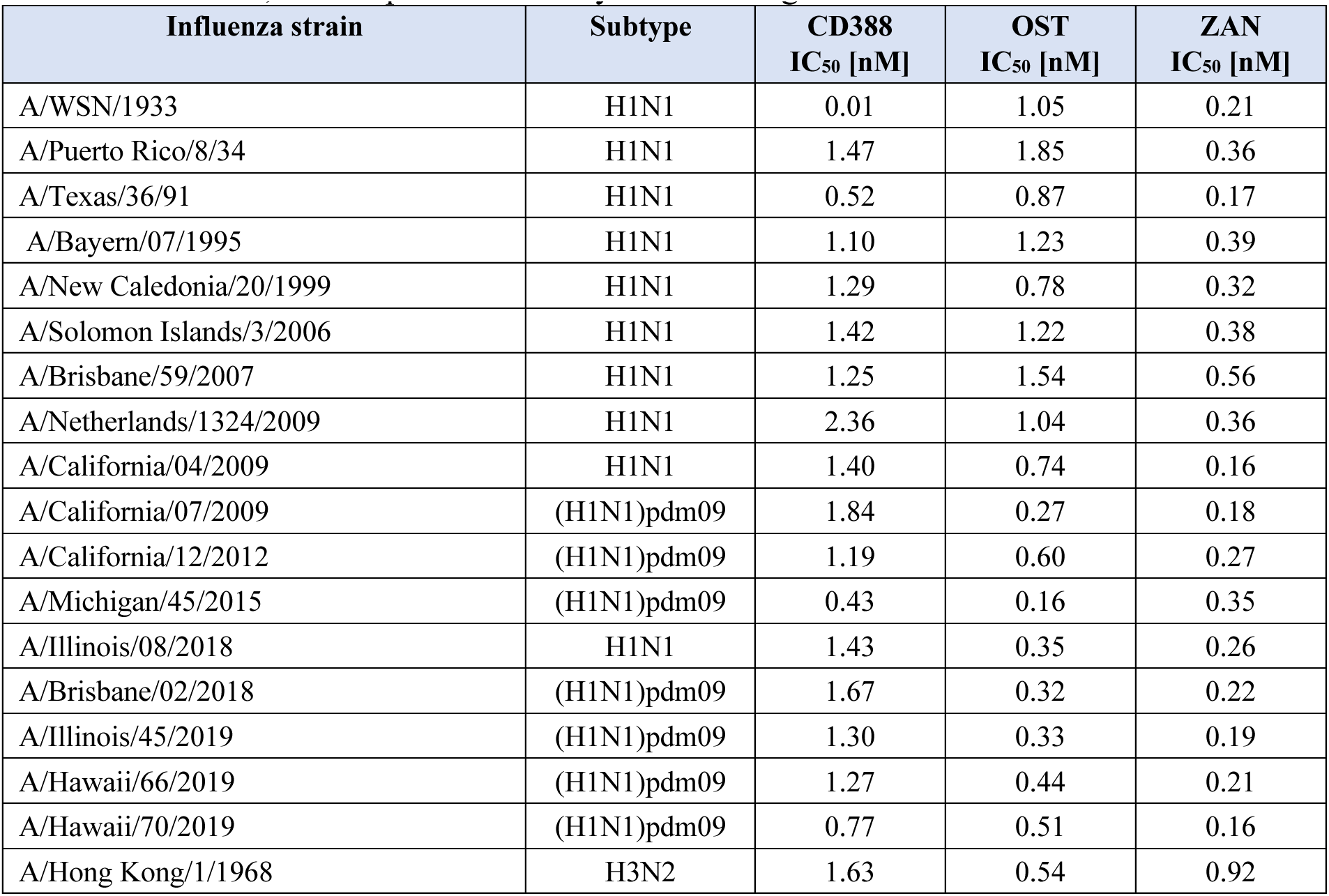
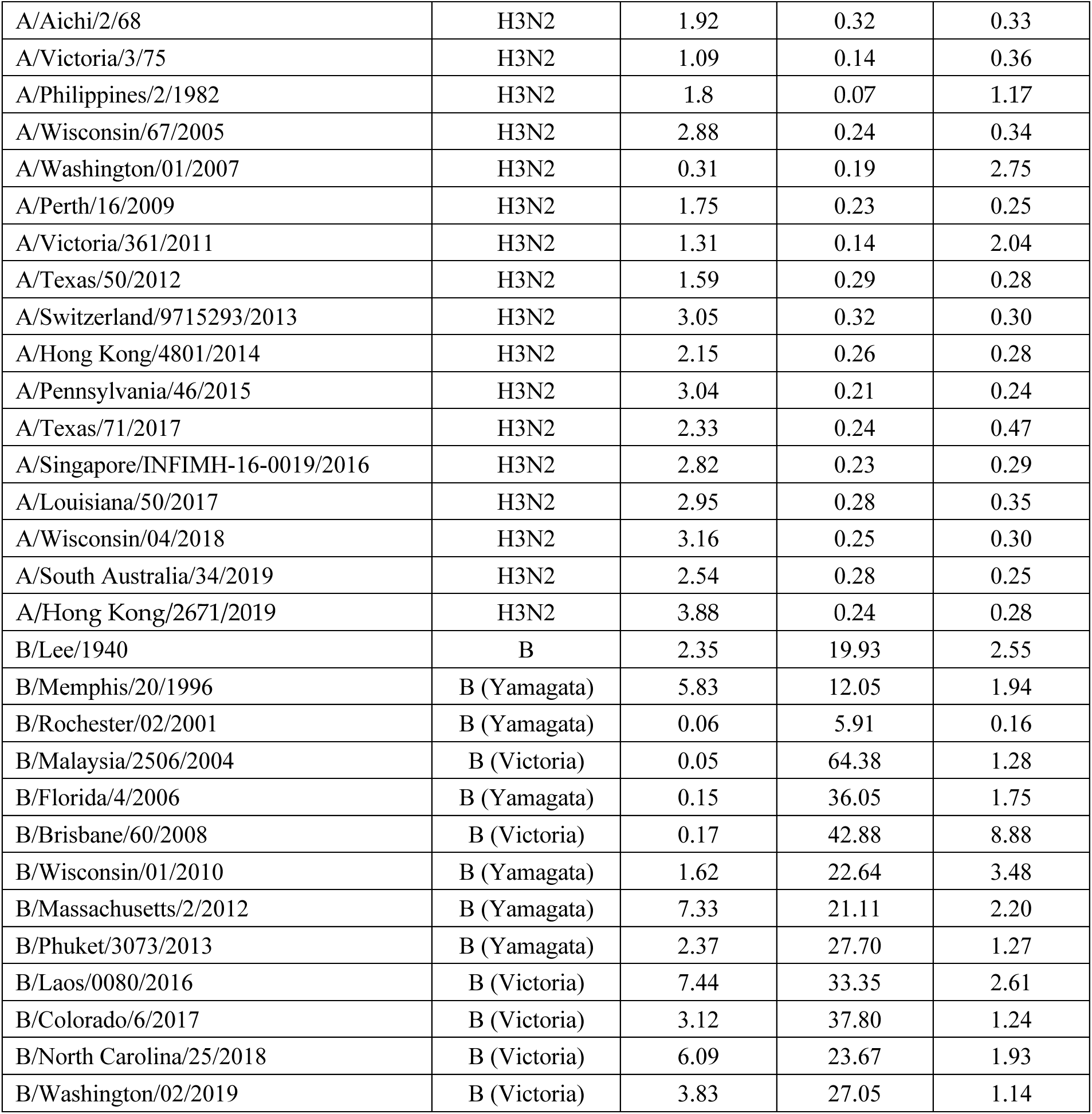
Universal, broad-spectrum activity of CD388 against influenza A and B in NA inhibition.

**Table 2.** Activity of CD388 against influenza A/H5N1 and A/H7N9 in NA inhibition (IC_50_)

| Influenza strain | Subtype | CD388<br>IC<sub>50</sub> [nM] | OST<br>IC<sub>50</sub> [nM] | ZAN<br>IC<sub>50</sub> [nM] |
| --- | --- | --- | --- | --- |
| A/Anhui/1/2005 | H5N1 | 3.72 | 2.96 | 0.25 |
| A/Anhui/1/2013 | H7N9 | 1.43 | 0.61 | 0.90 |
| A/Shanghai/1/2013 | H7N9 | 1.03 | >1,000 | 116.00 |

Greater discrimination between CD388 and ZAN or OST was observed in cell-based CPE assays. CD388 exhibited potent activity ranging from 0.01 nM to 11.25 nM against a large panel of influenza A and B viruses. CD388 demonstrated a median EC_50_ of 0.80 nM (*n* = 15; ranging from 0.01 to 11.25 nM) against subtype A/H1N1, 1.27 nM (*n* = 16; ranging from 0.03 to 8.53 nM) against subtype A/H3N2, and 1.72 nM (*n* = 13; ranging from 0.03 to 8.71 nM) against influenza B (**Fig. 1b-d**; **Extended Data Table 1**). Remarkably, the median EC_50_ for CD388 was >2 logs lower than for OST and ZAN against the panels of influenza A and B viruses. Similar enhancements in antiviral activity with sufficiently spaced multimeric NAI constructs compared to their monomeric counterparts has been observed previously^28,29^. Improved activity was attributed to inhibitor-mediated viral aggregation in solution and increased viral immobilization on the surface of infected cells as confirmed by electron microscopy^28,29^. When CD388 was tested against HPAI strains in cell-based microneutralization assays, median EC_50_ values were 0.57 nM versus H5N1 strains (*n* = 4; ranging from 0.09 to 0.95 nM) and 0.04 nM versus an A/H7N9 strain (*n* = 1) showing improved activity compared to OST and ZAN (**Fig. 1 e, f**; **Extended Data Table 2**). CD388 demonstrated comparable activity to baloxavir acid (BXA, the active moiety of BXM), against most influenza A and B strains tested (**Fig. 1b-d**; **Extended Data Table 1**). BXA was exceptionally potent against HPAI strains, with EC_50_ values below the lowest tested concentration of 0.03 nM (**Fig. 1 e, f**; **Extended Data Table 2**). Finally, the CC_50_ for CD388 in MDCK-SIAT1 cells was >10,000 nM (**Extended Data Tables 3, 4**), resulting in a >1,000-fold selectivity index with CD388 for influenza A and B in cell-based assays (**Extended Data Table 5**). Collectively, these data demonstrate that CD388 possesses highly improved activity and spectrum compared with small molecule NAIs in cell-based assays.

**Table 3.** NA inhibition (IC_50_) by CD388 vs other NAIs against CDC NA inhibitor susceptibility reference virus panel (versions 2.0 and 3.0) and the A/Bethesda/956/2006 (H3N2) NA R292K variant.

| Influenza strain | NA Genotype | CD388 |  | OST |  | ZAN |  |
| --- | --- | --- | --- | --- | --- | --- | --- |
|  |  | IC<sub>50</sub> [nM] | Fold change | IC<sub>50</sub> [nM] | Fold change | IC<sub>50</sub> [nM] | Fold change* |
| A/California/12/2012 (H1N1)pdm09 | H275 | 1.48 | -- | 0.99 | -- | 0.42 | -- |
| A/Texas/23/2012 (H1N1)pdm09 | H275Y | 1.96 | 1.3 | 633.4 | 639.8 | 0.56 | 1.3 |
| A/Illinois/45/2019 (H1N1)pdm09 | H275 | 1.30 | -- | 0.3 | -- | 0.19 | -- |
| A/Alabama/03/2020 (H1N1)pdm09 | H275Y | 0.98 | 0.76 | 426.8 | 1303.6 | 0.16 | 0.83 |
| A/Bethesda/956/2006 (H3N2) | R292K | 3.09 | n/a | >1,000 | n/a | 8.46 | n/a |
| A/Washington/01/2007 (H3N2) | E119 | 0.43 | -- | 0.29 | -- | 4.51 | -- |
| A/Texas/12/2007 (H3N2) | E119V | 1.95 | 4.5 | 179.6 | 619.3 | 1.59 | 0.4 |
| A/Pennsylvania/46/2015 (H3N2) | E119 | 3.04 | -- | 0.21 | -- | 0.24 | -- |
| A/Washington/33/2014 (H3N2) | E119V | 4.09 | 1.3 | 49.9 | 232.6 | 0.53 | 2.23 |
| B/Rochester/02/2001 | D198 | 0.12 | -- | 38.5 | -- | 1.15 | -- |
| B/Rochester/02/2001 | D198N | 0.65 | 5.4 | 291.5 | 7.6 | 14.02 | 12.2 |
| B/Memphis/20/1996 | R152 | 7.63 | -- | 15.61 | -- | 2.95 | -- |
| B/Memphis/20/1996 | R152K | 6.71 | 0.9 | >1,000 | >64.1 | 109.4 | 37.1 |
| B/North Carolina/25/2018 | D197 | 6.09 | -- | 23.67 | -- | 1.93 | -- |
| B/Missouri/12/2018 | D197E | 6.19 | 1.0 | 197.4 | 8.34 | 14.21 | 7.35 |
| B/Laos/0080/2016 | H134 | 7.44 | -- | 33.35 | -- | 2.61 | -- |
| B/Laos/0654/2016 | H134N | 4.66 | 0.6 | 171.8 | 5.15 | 310.80 | 119.3 |

n/a = not available; \*Fold-change in IC<sub>50</sub> compared to parental strain.

### CD388 is efficacious in mouse models

The efficacy of CD388 in lethal mouse infection models was evaluated. CD388 administered as a single IM dose 2 h post-challenge was fully protective at 0.3 mg/kg against a lethal challenge with mouse-adapted influenza A/Puerto Rico/8/1934 (H1N1) (**Fig. 2a**). Mice treated with unconjugated Fc or PBS as vehicle control succumbed to infection at similar times post-challenge. Following treatment with CD388 at the minimal protective dose (0.3 mg/kg), only limited, transient body weight loss occurred (**Fig. 2b**). To better understand the mechanism for CD388-mediated protection, the viral load and cytokine levels in lungs were determined four days after infection in this model. CD388 demonstrated a dose-dependent reduction in plaque-forming units [PFU]/g in lung tissue of 0.83 logs at 0.03 mg/kg to below the limit of detection (>3 logs) at 3 mg/kg by plaque assay (**Fig. 2c**). Additionally, CD388 showed a dose-dependent reduction in lung pro-inflammatory cytokines IL-6 and MCP-1. At the 3 mg/kg dose of CD388, cytokine levels declined to those present in uninfected control mice lungs (**Fig. 2d, e**). Interestingly, OST administered BID x 5 days at the human equivalent dose (5 mg/kg) in the same model was not protective, although it delayed time of death (**Fig. 2f**). OST administered at 10x the human equivalent dose (50 mg/kg) resulted in 80% survival (**Fig. 2f**) but that benefit was accompanied with substantial body weight loss in surviving mice (**Fig. 2g**). A dose response in viral burden reduction was not observed with OST administered at 5 mg/kg versus 50 mg/kg (BID for 4 days) and resulted in minimal, non-dose dependent viral load reductions of ca. 0.66 log or 0.09 logs, respectively (**Fig. 2h**). However, OST at both doses significantly reduced lung IL-6 and MCP-1 cytokine levels (**Fig. 2i, j**). In conclusion, CD388 demonstrated significantly improved efficacy compared to OST in this model.

**Figure 2.**
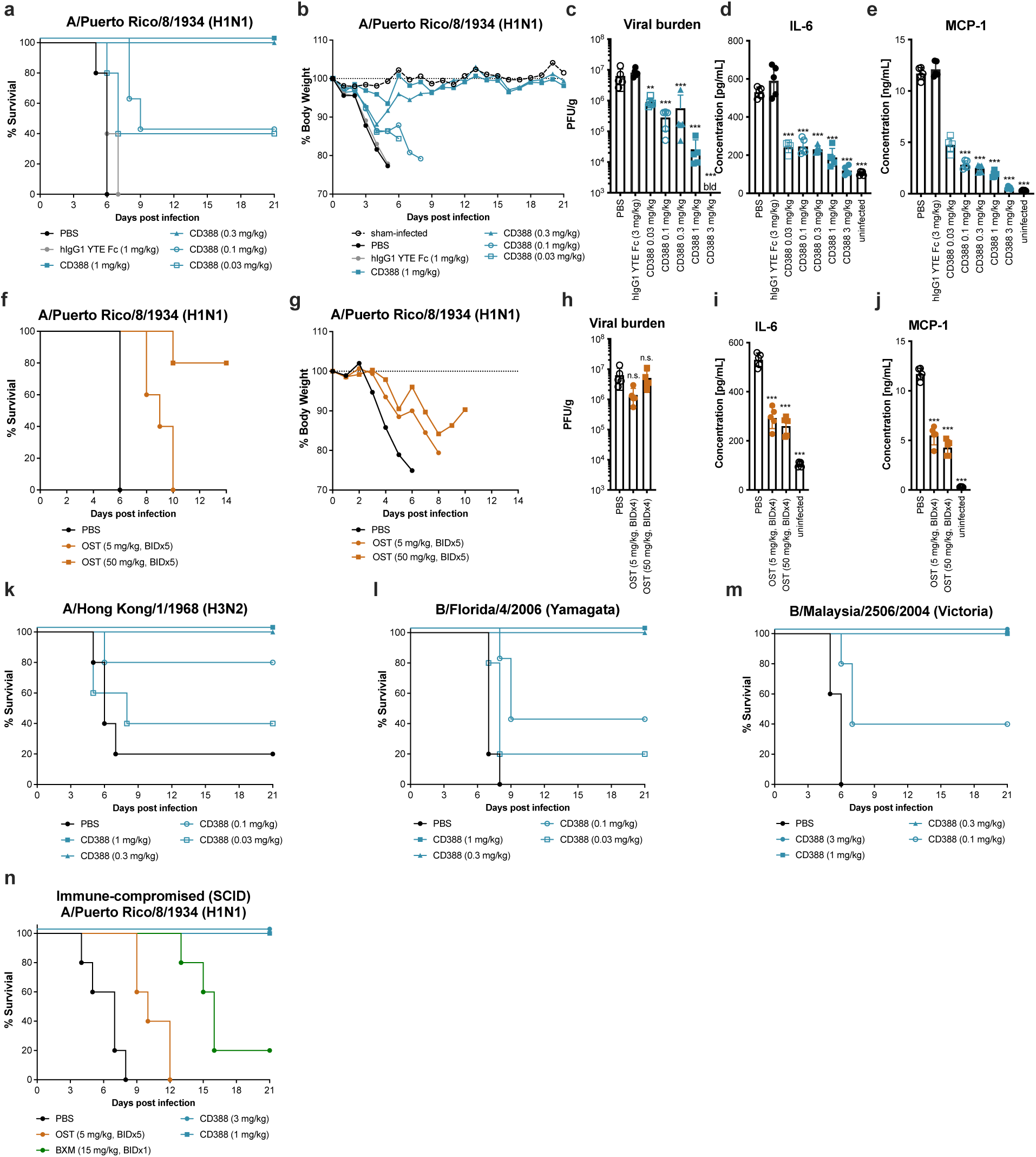
Universal efficacy of CD388 in lethal mouse models of influenza A and B infection. Efficacy of CD388 dose response on (**a**) survival and (**b**) % body weight change and the effect of CD388 on (**c**) lung viral burden (mean ± SD; *n* = 5/group) and pro-inflammatory lung cytokine levels of (**d**) IL-6 and (**e**) MCP-1 at 4 days post-infection following lethal challenge with mouse-adapted influenza A/Puerto Rico/8/1934 (H1N1) in BALB/c mice. Efficacy of OST on (**f**) survival and (**g**) % body weight change in a lethal mouse model with mouse-adapted influenza A/Puerto Rico/8/1934 (H1N1) in BALB/c mice (*n* = 5/group) and effect of OST on (**h**) lung viral burden and pro-inflammatory lung cytokine levels of (**i**) IL-6 and (**j**) MCP-1 at 4 days post-infection. Efficacy of CD388 in lethal mouse models of (**k**) A/Hong Kong/1/1968 (H3N2), (**l**) B/Florida/4/2006 (Yamagata) and (**m**) B/Malaysia/2506/2004 (Victoria). (**n**) Efficacy of CD388 against mouse-adapted A/Puerto Rico/8/1934 (H1N1) in immune-compromised BALB/c SCID mice (*n* = 5/group). Statistical analyses were performed using one-way ANOVA with Dunnett’s multiple comparison test (*P<0.05, ** P<0.01, ***P<0.001; ns, not statistically significant) in GraphPad Prism version 9 software.

In vitro activity and spectrum of CD388 also translated to efficacy against representative viruses belonging to the A/H1N1 and A/H3N2 subtypes, as well as the two influenza B lineages Victoria and Yamagata, in lethal mouse models. Single IM doses of 0.3 mg/kg of CD388 were fully protective against A/Hong Kong/1/1968 (H3N2), B/Florida/4/2006 (Yamagata) and B/Malaysia/2506/2004 (Victoria) (**Fig. 2k-m**; **Extended Data Fig. 2a-c**). CD388 demonstrated efficacy following a single IM dose at 1 mg/kg or lower against a larger panel of mouse-virulent influenza strains including A/WSN/1933 (H1N1), A/California/07/2009 (H1N1)pdm, A/California/12/2012 (H1N1)pdm09, A/North Carolina/04/2014 (H1N1)pdm09, A/Hawaii/70/2019 (H1N1)pdm09, and B/Colorado/6/2017 (Victoria) (**Extended Data Fig. 2d-i**; **Extended Data Table 6**). Similar to the results observed with A/Puerto Rico/8/1934 (H1N1), CD388 treatment at the minimal protective dose corresponded with only minimal, transient body weight loss with the other strains tested (**Extended Data Fig. 2**). To assess the potential of CD388 to prevent infections in high-risk groups including the elderly and immune-compromised, we evaluated the CD388 efficacy in severe combined immune-deficiency (SCID) mice, which are deficient in T cells and B cells. CD388 demonstrated full protection with a single 1 mg/kg dose administered IM in SCID mice infected with a lethal challenge of A/Puerto Rico/8/1934 (H1N1), whereas OST and BXM were ineffective or only partially protective (20%), respectively (**Fig. 2n**). Taken together, these data highlight the potential of CD388 to provide universal influenza protection in healthy and high-risk populations with similar doses.

### CD388 is efficacious in prophylaxis

Pharmacokinetic (PK) profiling and prophylactic efficacy studies were conducted to evaluate the potential of CD388 for use as a durable, long-acting agent for universal influenza prevention. The half-life of CD388 after a single IV dose in mice or cynomolgus monkeys was 106 h and 364 h following 1- or 8-week PK studies, respectively (**Extended Data Table 7**). As expected, the engineered YTE-Fc used in CD388 demonstrated improved binding compared to unconjugated human wild-type (WT) Fc to human FcRn (in a pH-dependent manner) by Biolayer interferometry (BLI) (**Extended Data Table 8**). CD388 exhibited dose-linear PK in mice across a wide IM dose range (0.3 – 30 mg/kg), with AUCs ranging from 107 to 6,507 h x µg/mL (**Fig. 3a**; **Extended Data Table 9).** The in vivo stability of intact CD388 was measured by comparing exposures using two different ELISA capture methods; one that measured Fc-levels with two distinct human Fc-capture antibodies, and one that measured intact CD388 using recombinant viral NA for capture and an anti-human Fc capture antibody for detection (**Extended Data Fig. 3**). CD388 plasma levels measured by both methods were comparable within experimental error, demonstrating that CD388 circulates as an intact conjugate in mice and monkeys. CD388 reached peak lung levels within 4 – 24 h after IM administration, allowing rapid onset of protection. CD388 dosed IM at 10 mg/kg rapidly distributed to epithelial lining fluid (ELF) at 34% of plasma levels (**Fig. 3b)**.

**Figure 3.**
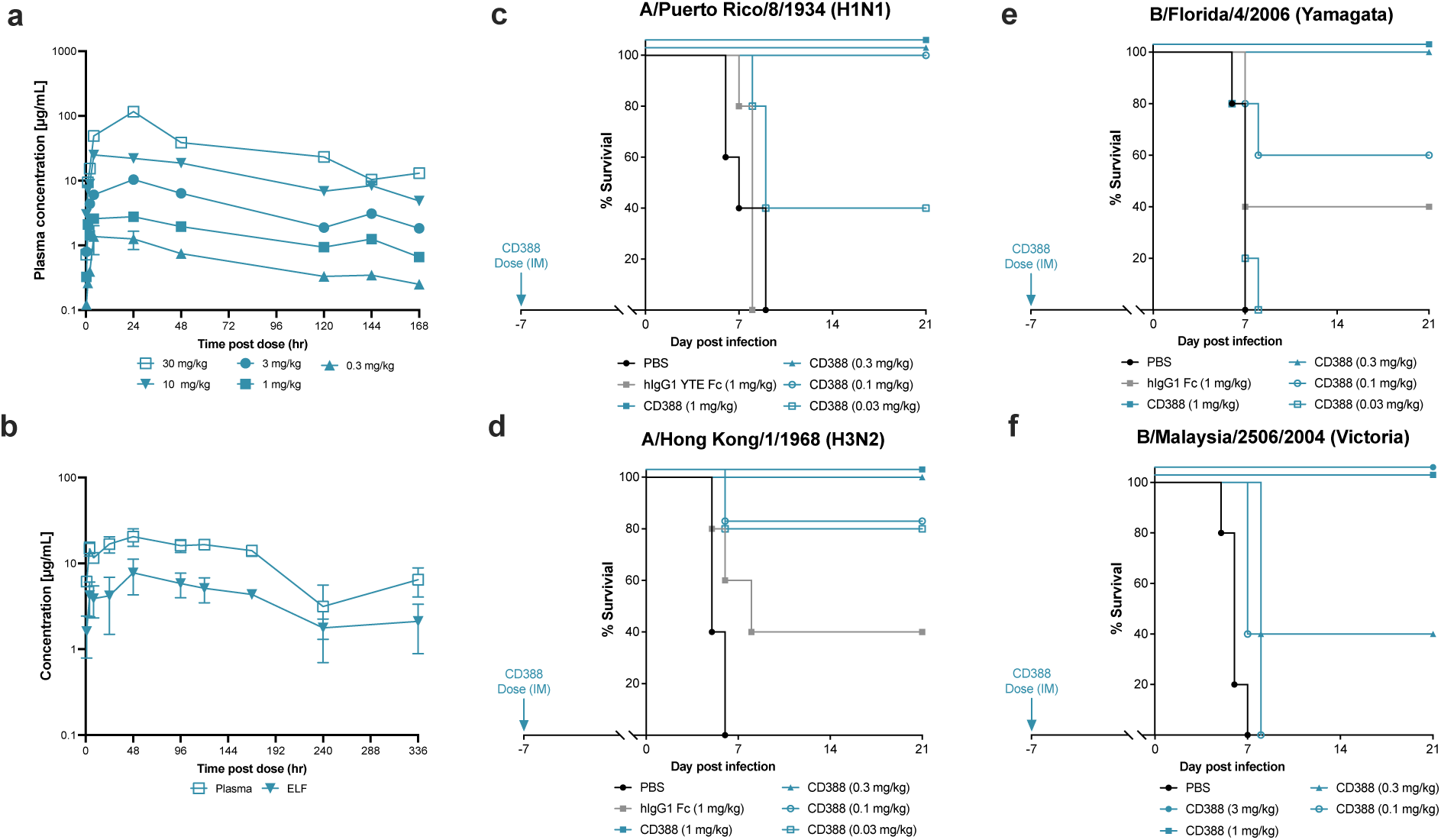
Mouse PK and prophylactic efficacy of CD388 in lethal mouse models of influenza A and B infection. (**a**) Plasma levels of CD388 at doses from 0.3 – 30 mg/kg following IM administration in mice (mean ± SD; *n* = 2/group) by NA capture. (**b**) CD388 levels in plasma and lung epithelial lining fluid (ELF) of mice after IM dosing (mean ± SD; *n* = 6/group) by NA capture. Protection provided by CD388 dosed as indicated IM 7 days prior to lethal challenge against influenza (**c**) A/Puerto Rico/8/1934 (H1N1), (**d**) A/Hong Kong/1/1968 (H3N2), (**e**) B/Florida/4/2006 (Yamagata), and (**f**) B/Malaysia/2506/2004 (Victoria) in BALB/c mice (*n* = 5/group).

To assess CD388 efficacy in a prevention setting, CD388 was administered to mice 7 days prior to infection. Single IM doses of 1 mg/kg conferred full protection against influenza A/Puerto Rico/8/1934 (H1N1), A/Hong Kong/1/1968 (H3N2), B/Florida/4/2006 (Yamagata), B/Malaysia/2506/2004 (Victoria) (**Fig. 3c-f**; **Extended Data Fig. 4a-d**; **Extended Data Table 6**), and numerous other mouse-virulent influenza strains (**Extended Data Fig. 5**). These data highlight the promise of CD388 as a therapeutic for universal prevention of seasonal and pandemic influenza.

### CD388 is active against NAI-resistant variants

The multivalent presentation of ZAN dimers in CD388 prevented loss in activity against viruses harboring resistance mutations to small molecule NAIs. Specifically, CD388 retained potent activity against the CDC NA inhibitor susceptibility reference virus panel (versions 2.0 and 3.0) comprising clinically relevant variants with reduced or highly reduced activity to NAIs, including H275Y (A/H1N1), E119V (A/H3N2), D197E (B), D198N (B), R152K (B), H134N (B), and the clinical isolate R292K (A/H3N2). While OST and ZAN lost more than 100-fold in potency against some of the highly resistant variants (**Table 3**), the IC_50_ values for CD388 were not significantly different against any of these variants when compared to parental strains. Next, the efficacies of OST and CD388 were evaluated against the OST-resistant NA variant H275Y in a lethal mouse model. The minimal protective dose of CD388 was 0.3 mg/kg, in line with protective dose levels against NA-sensitive viruses in lethal mouse models (**Extended Data Table 6**). OST dosed at the human equivalent dose of 5 mg/kg BID for 5 days did not confer any survival benefit (**Fig. 4a**). Similarly, the efficacies of CD388 and ZAN were compared in lethal challenge models with paired ZAN-sensitive and ZAN-resistant NA H134N strains. CD388 demonstrated identical minimal protective doses of 0.3 mg/kg against both strains (**Fig. 4b, c**), while the minimal protective dose of ZAN shifted from 1 mg/kg against the ZAN-sensitive virus to 10 mg/kg against the ZAN-resistant variant (**Fig. 4d, e**). The retention of potency observed with CD388 against clinically observed NA mutants with reduced susceptibility to all other approved small molecule NAIs highlights the potential durability of CD388 as a preventative agent against seasonal and pandemic influenza.

**Figure 4.**
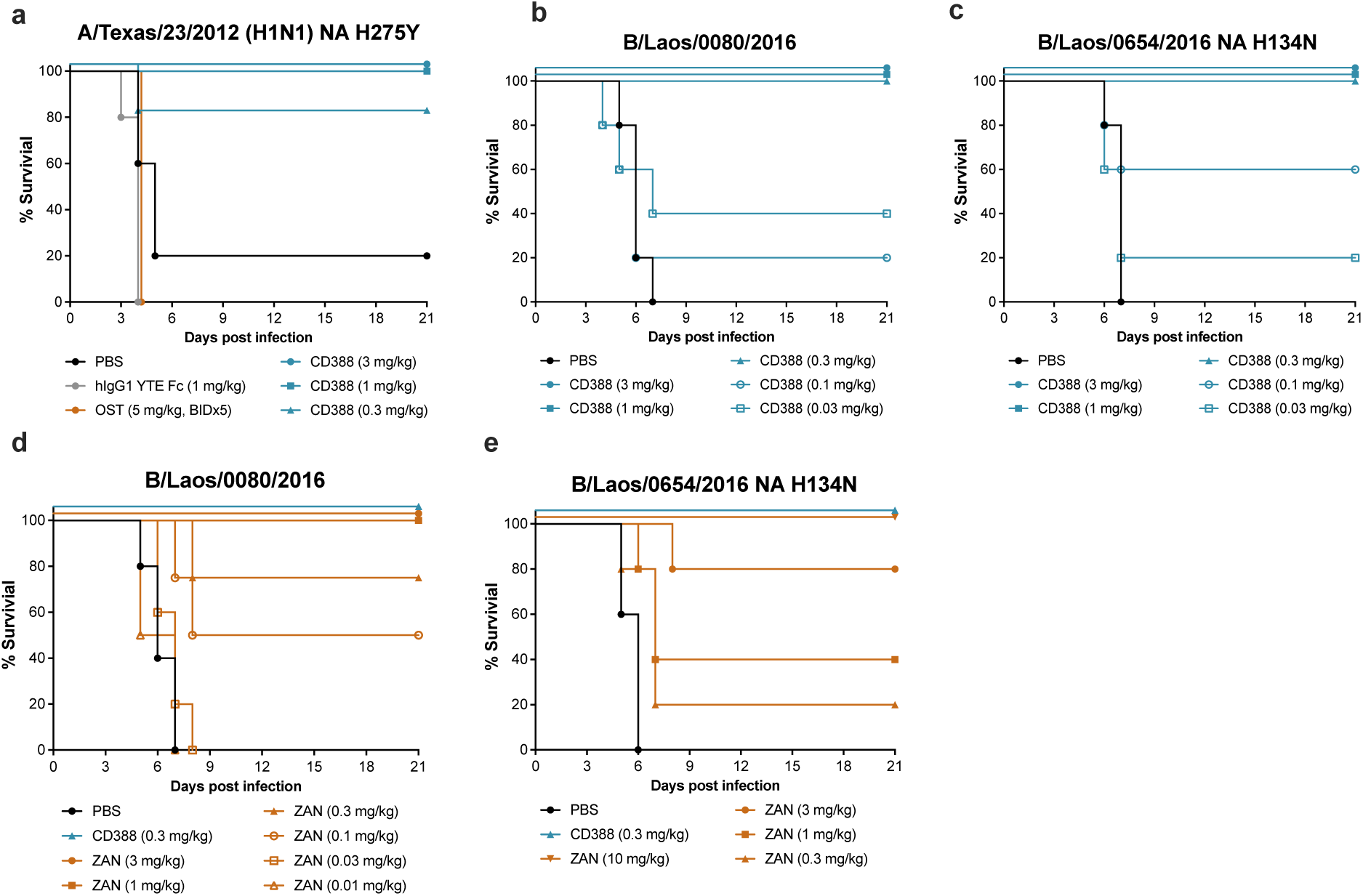
CD388 efficacy in lethal mouse models of influenza A and B harboring NA mutations that confer reduced susceptibility to other NAIs. Efficacy of CD388 administered IM at the indicated doses at 2 h post-infection against (**a**) influenza A/Texas/23/2012 (H1N1)pdm09 NA H275Y variant, (**b**) ZAN-susceptible influenza B/Laos/0080/2016 (Victoria) and (**c**) ZAN-resistant B/Laos/0654/2016 NA H134N variant (Victoria) in BALB/c mice (*n* = 5/group). Comparisons of dose-responses of ZAN administered intranasally starting at 2 h post-infection for 5 days and CD388 administered IM once 2 h post-infection at 0.3 mg/kg to ZAN-susceptible influenza (**d**) B/Laos/0080/2016 and (**e**) ZAN-resistant B/Laos/0654/2016 NA H134N variant in BALB/c mice (*n* = 5/group).

### CD388 shows low potential for development of resistant influenza variants

To assess the potential of influenza resistance development against CD388, serial passage experiments were conducted in MDCK cells infected with representative influenza A/H1N1 (n=2), A/H3N2 (n=2) and B (n=2) viruses utilizing two drug selection methodologies: static, sub-inhibitory and incremental dose-escalation. Serial passage of influenza A and B in the presence of CD388 resulted in minimal changes in the susceptibility of all strains evaluated when comparing plaque-purified viruses isolated following 10 rounds of passaging to their isogenic parental strains. Genotypic analyses identified a single NA residue (S247 or A246 using N1 or N2 numbering, respectively) that was substituted to arginine or valine following prolonged exposure CD388. This residue is proximal to the CD388/NA binding site and mutations were identified in representative strains of A/H1N1 (S247R; S231R based on A/WSN/1933 strain numbering) and A/H3N2 (A246V, A/Victoria/3/75). Both NA amino acid substitutions have been documented previously^32,33^ and occur at very low frequencies amongst clinical isolates.

We assessed the cross-resistance of the above identified S247R; S231R NA variants in cell-based plaque reduction assay (PRA). CD388 lost less than 3-fold activity in PRA. Thus, compared to other NAIs, CD388 demonstrated equivalent or higher potency (**Fig. 5a, b**; **Extended Data Table 10**). Importantly, the EC_50_ values for CD388 against these variants were still within the median EC_50_ ranges observed against the respective susceptible subtypes. A low predisposition of influenza A and B to develop reduced susceptibility to CD388, lack of novel mutations recovered, and equivalent or higher levels of reduced susceptibility of approved NAIs to the selected resistance substitutions at NA residue A246/S247, suggest a low potential for clinical resistance development that is consistent with that of ZAN.

**Figure 5.**
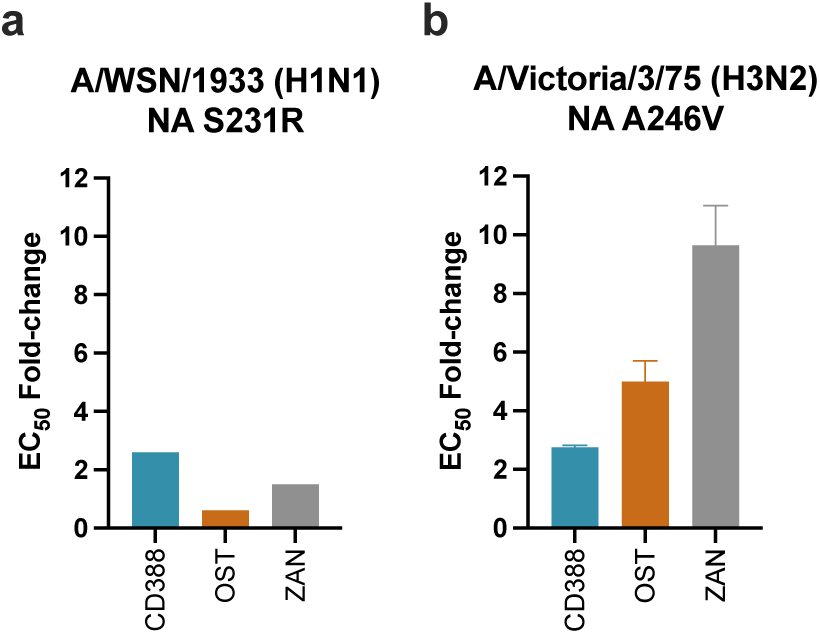
Cross-resistance of CD388-selected NA variants to OST and ZAN. Fold-change of a plaque-purified clone isolated at passage 10 versus parental strains by EC_50_ [nM] to CD388-selected NA variants (**a**) S231R in A/WSN/1933 (H1N1) or (**b**) NA A246V in combination with N113H/T453I in A/Victoria/3/75 (H3N2) for CD388, OST, and ZAN in cell-based PRA.

## DISCUSSION

Seasonal influenza continues to drive an unacceptably high incidence of severe illness and mortality, with the potential for even worse outcomes when novel, highly pathogenic, zoonotic strains become transmissible to humans, currently an issue of significant concern with the emergence of H5N1 influenza in wild and domestic animal populations. After two influenza seasons during which influenza infections were reduced by SARS-CoV-2 pandemic precautions, influenza infections rebounded in the early 2022-23 season, and now continue at pre-pandemic levels. From October 1, 2023 through May 18, 2024, the CDC estimates there have been 35-64 million influenza infections in the US, leading to 390,000 – 820,000 hospitalizations and 25,000 - 71,000 deaths^34^. Limited vaccination coverage in the US (ca. 50%), limited VE (10-60% depending on season) and poor immune responses to vaccines in immune-senescent populations leave large segments of the human population poorly protected from influenza and at higher risk of severe infections or death^35^. Additionally, waning immunity to influenza has been shown to result in reduced overall VE within a single influenza season^36,37^. Next-generation vaccines or novel therapeutic agents with broad-spectrum coverage are needed to reduce the number of severe cases, hospitalization, deaths, and the economic burden due to seasonal influenza^38^.

In recent years, mAbs have emerged as promising strategies for treatment and prevention of viral diseases, including influenza^39^. Antibodies raised against HA and NA have been shown to correlate with protection^40,41^. However, most efforts to develop mAbs against influenza have focused on the conserved stalk domain of HA. To date, no HA antibodies have successfully advanced in clinical trials, and none provided coverage against influenza B, which is responsible for, on average, 25% of influenza infections^42^. Notably, in the early 2019-20 season, influenza B viruses dominated, and were responsible for nearly 60% of influenza cases in the US^43^ underscoring the need for universal protection against influenza A and B.

NA has long been recognized as an attractive alternative target to HA for development of anti-influenza therapies that will provide broader coverage^4,10^. This assertion is supported by a recent human challenge study that demonstrated a superior correlation between reduced disease severity and the detection of high anti-NA titres compared to high anti-HA titres^44^. More recently, anti-NA mAbs isolated from patients infected with influenza were described that have broader coverage compared to anti-HA mAbs with fewer spectrum gaps^14,45,46^. Nonetheless, the best success in expanding the spectrum of coverage has been achieved by targeting the strictly conserved NA active site with small molecules. As a stable, multivalent conjugate of a novel ZAN dimer with an extended Fc domain of a human IgG1 antibody engineered to prolong half-life, CD388 leverages and improves upon the desirable attributes of potent, broad-spectrum small molecule NAIs and long acting mAb therapeutics. ZAN was selected for conjugation due to its best-in-class antiviral potency and resistance profile compared with other NAIs. The multivalent presentation of ZAN dimers on CD388 further improves its activity and breadth of coverage to encompass clinically observed NAI-resistant influenza variants. The low resistance potential of CD388 presents a conceivably more durable profile than small molecule NAIs, vaccines, and mAb-based options. Conjugation to the engineered Fc domain results in a long half-life, similar to that of mAbs, with distribution to lung that is superior to what has been reported for mAbs^47^. These features bestowed exceptionally potent efficacy in prevention and treatment to CD388, with dose-dependent reductions in lung viral burden and superior activity compared to OST and ZAN. Importantly, CD388 efficacy was retained in an immune-compromised mouse model, suggesting that efficacy is driven predominantly by the intrinsic antiviral activity of CD388, with minimal reliance on Fc-mediated immune effector function. Taken together, the preclinical data reported here highlight the potential for use of CD388 as a universal, long-acting therapeutic for the treatment and prevention of seasonal and pandemic influenza in both healthy and high-risk individuals. CD388 has advanced to clinical development, where it will be evaluated in a Phase 2b study for universal prevention of influenza during the 2024-2025 flu season in the Northern Hemisphere.

## ACKNOWLEDGEMENTS

The authors would like to thank Dr. Ken Bartizal (Cidara Therapeutics), Paul De Jesus, Dr. Laura Martin-Sancho, Dr. Sumit Chanda (Scripps Research, CA) and Dr. Stacey Schultz-Cherry (St. Jude Children’s Research Hospital, TN) for their assistance, expert insight and for kindly providing reagents. The authors are also grateful to Dr. Bonnie Bassler (Princeton, NJ) for her critical review of this manuscript and editorial suggestions.

## AUTHOR INFORMATION

These authors contributed equally: Simon Döhrmann, James Levin, Jason N. Cole, Allen Borchardt

## Contributions

Experiments were designed and conceived by S.D., J.L., J.N.C, J.L., V.O. and L.W.T. CD388 was invented and synthesised by A.B., Z.Y.C., T.L., T.P.B. and L.W.T. Protein production was done by J.F. In vitro experiments were performed by S.D., J.N.C., A.A., E.A., R.G., D.Z. and N.D. Peptide mapping experiments and analysis were conducted by G.H. The in vivo studies were designed by J.L. and carried out by K.A., S.D., A.A., E.A. Figures were prepared by S.D., J.L. and L.W.T. The manuscript was written by S.D., J.L., J.N.C. and L.W.T. All authors contributed to editing and reviewing the manuscript and support the conclusions.

## DATA AVAILABILITY STATEMENT

The authors declare that all relevant data supporting the key findings of this study are available within the paper and its supplementary information. Source files are available from the corresponding author upon reasonable request.

**SUPPLEMENTARY INFORMATION** is available for this paper.

## METHODS

### Viral strains

Influenza viruses were obtained through BEI Resources (Manassas, VA), NIAID, NIH, the International Reagent Resource (Manassas, VA), CDC’s WHO Collaborating Center for Surveillance, Epidemiology and Control of Influenza (Atlanta, GA); and Microbiologics (San Diego, CA). Influenza strains A/WSN/1933 (H1N1) and B/Malaysia/2506/2004 (Victoria) were kindly provided by Dr. Sumit Chanda (Scripps Research, CA). A complete list of viruses used herein is shown in **Supplemental** Table 1. Influenza strains were propagated in Madin Darby Canine Kidney (MDCK) or MDCK SIAT1 (Sigma, 05071502) at 37 °C, 5% CO_2_ until 80% of CPE. Cell-free supernatant was obtained by centrifugation at 500 × *g* for 5 min and virus was stored at −80 °C. PFU were determined by plaque assay. Briefly, 100 µL of virus dilutions were added to confluent monolayer of MDCK cells in 24-well plates and incubated for 1 h at room temperature with rocking every 15 min. After removing the virus, liquid overlay media containing Avicel was added to MDCK cells. Cells were incubated at 37°C, 5% CO_2_ for 40 – 72 hours. After incubation and media removal, cells were stained with crystal violet to enumerate plaques.

### Recombinant NA

Recombinant NA from cell lysate of A/H5N1 and AH7N9 viruses (Sino Biological, TX) are summarized in **Supplemental** Table 2.

### NA inhibition

CD388, OST or ZAN were tested at 0.001 – 1000 nM (25 µL/well) in 96-well plates (Corning). Influenza virus or NA from cell lysate (Sino Biological) was diluted to be within the linear range of detection. Virus and test article were incubated for 20 min prior to the addition of NA substrate at 37°C, 5% CO_2_. NA inhibition activity was determined using the commercial NA-Fluor kit (Applied Biosystems) according to the manufacturer’s instructions. Fluorescence was measured with the EnSpire multimode plate reader (PerkinElmer). IC_50_ values were calculated with GraphPad Prism version 8 using nonlinear regression analysis (dose-response inhibition).

### CPE

MDCK or MDCK SIAT1 cells were seeded in 96-well plates at 1-2 × 10^4^ cells/well in DMEM (Gibco) supplemented with 10% FBS (Gibco). Cells were incubated for 18-24 h at 37°C, 5% CO_2_. The next day, the media of a confluent cell monolayer (>90%) was aspirated and washed with PBS. CD388, OST, ZAN or BXA were tested at concentrations ranging from 0.001 pM to 10,000 nM. Subsequently, cells were infected with influenza A or B virus in DMEM in the presence of 1 µg/mL TPCK-Trypsin to yield >70% CPE in the untreated control. CPE was determined after fixation and staining with 0.1% crystal violet after 3 days for influenza A and after 5 days for influenza B. The absorbance was read at 595 nm on an EnSpire multimode plate reader. EC_50_ values were calculated with GraphPad Prism version 8 using nonlinear regression analysis (dose-response inhibition).

### PRA

PRA were performed on starting and final passage viruses in MDCK or MDCK-SIAT1 cells for sub-inhibitory H3N2 passaged viruses and cells were seeded in 24-well plates at 5 × 10^5^ cells/well in 0.5 mL of DMEM containing 10% FBS and incubated for approximately 24 h at 37°C, 5% CO_2_. CD388, OST and ZAN were diluted in infection buffer containing PBS, 0.35% BSA, 1.105 mM CaCl_2_ and 2.24 mM MgCl_2_. Test articles were pre-incubated with virus for 30 min at room temperature prior to addition to cell monolayers. The multiplicity of infection (MOI) for each drug-virus combination was selected to produce 30 plaques in the PBS control well. Adsorption was carried out for 1 h, the virus-test article mix was removed, and the infected cells were incubated for 48 h in the presence of test article diluted in a mixture of 1.25% Avicel, DMEM, 0.01% DEAE-dextran and 2 μg/mL of TPCK trypsin at 37°C, 5% CO_2_. After 48 h the Avicel mixture was then removed, and cells were fixed with paraformaldehyde and stained with 1% crystal violet to count the plaques. EC_50_ values were calculated using GraphPad Prism version 8.

### Microneutralization

This method was performed as described by Viroclinics (Rotterdam, Netherlands)^48^. Briefly, a twofold dilution series of CD388, OST, ZAN or BXA was performed and mixed with virus aiming for 50-100 PFU/well. The virus-test article mixture was immediately added to MDCK cells and incubated for 90 min at 37°C and 5% CO_2_. After incubation, the inoculum was removed, cells were washed and replaced with medium containing the same dilutions of test articles. The plates were incubated for ∼24 h at 37°C and 5% CO_2_. After completion of the incubation period, the cells were formalin-fixed and permeabilized with 70% ethanol. They were then incubated with a primary mouse antibody directed against the viral influenza A nucleoprotein (clone Hb65, EVL), followed by incubation with a secondary goat anti-mouse IgG HRP peroxidase conjugate. TrueBlue substrate added, resulting in the development of a blue precipitate on virus-positive cells. Images of all wells were acquired by a CTL Immunospot analyzer, equipped with software to quantitate the virus-positive cells (= virus signal). Quantification was done using ‘well area covered’ (WAC) counts. The EC_50_ was calculated according to the method as previously described^49^.

### PK ELISA

Nunc MaxiSorp 96-well plates were coated with 0.1 U/well NA from A/California/04/2009 (H1N1) (Sino Biological) for NA capture in 1x KPL coating buffer, or with 0.1 µg/well of mouse anti-human IgG (C_H_2 domain) clone R10Z8E9 (BioRad) for Fc capture in carbonate buffer. Plates were incubated at room temperature (RT) for 1 h on an orbital plate shaker at 500 rpm for NA capture or stationary at 4°C overnight for Fc capture. Plates were washed 5x with PBST and blocked with 1% BSA (MilliporeSigma) for NA capture or 5% non-fat dry milk in PBST (Cell Signalling) for Fc capture for 1 h at RT with shaking. NA sample diluent was PBS with 0.5% BSA, 0.025% Tween 20. Fc diluent was PBS with 2.5% non-fat dry milk and 0.025% Tween 20. Duplicate serial dilutions of the plasma samples were plated at 100 µL/well and incubated at RT for 2 h. Sample diluent was prepared with species-matched plasma (cynomolgus monkey at final dilution of 1:2,500; mouse at final dilution of 1:100). CD388 standard curves ranged from 0.230 to 500 ng/mL for NA capture and 0.03 – 55 ng/mL for Fc capture. Curves were run on each plate in duplicate. Following the 2 h incubation, plates were washed 5x with 300 µL/well PBST. Conjugate bound to NA or anti-Fc antibody on the plates was then probed with an HRP-conjugated anti-human IgG Fc F(ab’)_2_ (Jackson ImmunoResearch) diluted 1:2,000 in sample diluent for 1 h at RT with shaking. Plates were then washed 8x with 300 µL/well PBST and developed with 100 µL/well TMB substrate for 7-8 min. The reaction was stopped with 100 µL/well 1N H_2_SO_4_. Absorbance was read at 450 nm with an EnSpire multimode plate reader. Plasma samples were interpolated following nonlinear regression analysis (Sigmoidal, 4PL analysis) of the standard curves in GraphPad Prism version 8.

### Sub-inhibitory, static serial passage

Sub-inhibitory, static serial passage studies were conducted in MDCK cells infected with influenza A/WSN/1933 (H1N1), A/Washington/01/2007 (H3N2) or B/Malaysia/2506/2004 (Victoria) at MOI of 0.05 at 37°C, 5% CO_2_ for 24 h. The concentrations for each test article were optimized to result in a ∼2-log reduction in titer as compared to the vehicle control, while maintaining sufficient virus for subsequent passages. CD388 was tested at 0.003125 – 0.25 nM, BXA at 4 – 20 nM, and OST at 4 – 200 nM or with PBS as a control. Following the addition of test articles, cells were incubated for 24 h. Next, viral supernatants were collected and the viral titer was determined by plaque assay. Freshly seeded cells were infected with viral supernatant adjusted to MOI 0.05. This process was repeated for 10 passages.

### Dose escalation serial passage

Dose-escalation serial passage studies were conducted in MDCK cells infected with influenza A/Hawaii/70/2019 (H1N1)pdm09, A/Victoria/3/75 (H3N2), or B/Phuket/3073/2013 (Victoria) at an MOI of 0.01 – 0.05 and incubated at 37°C, 5% CO_2_ for 24 h (influenza A) or 48 h (influenza B). The concentrations for each test article were optimized to result in a ∼2-log reduction in titer as compared to the vehicle control, while maintaining sufficient virus for subsequent passages. CD388 was tested at 2 – 8 nM, BXA at 4 – 20 nM, and OST at 4 – 200 nM. A PBS control was also included. Following the addition of test articles, cells were incubated for 24 h. After 24 h, viral supernatants were collected, and the viral titers were determined by plaque assay. Freshly seeded cells were infected with viral supernatant adjusted to MOI 0.05. Each virus was serially passaged in the presence of 2-fold increasing concentrations of the test article until viruses could not be propagated further. The susceptibility of individual clones isolated from final passage was determined against test articles using a PRA.

### Sequencing of viral RNA

Viral RNA was extracted from influenza strains using a QIAamp Viral RNA Mini Kit (QIAGEN). To prepare samples for WGS, RNA was reverse transcribed and the entire genome of influenza was amplified in a single RT-PCR reaction using the Uni/Inf primer set^50^. PCR products were cleaned and concentrated using a DNA Clean and Concentrator-25 kit (Zymo Research). DNA concentrations were determined using a NanoDrop spectrophotometer (ThermoFisher) prior to WGS analysis. WGS was performed on individual starting and final passage plaque-purified strains at BATJ (San Diego, CA). Library construction and quantification for each sample started with ∼500 ng viral cDNAs that were then enzymatically fragmented to ∼200-500 bp pieces. NGS libraries were then constructed using the TruSeq Nano DNA Library Prep Kit (Illumina), which included steps of end-repairing, library size-selection, 3’-end adenylation, ligation of indexed adaptors, and PCR enrichment. The constructed libraries were then quantified with Qubit Fluorometer (ThermoFisher). Sequencing was performed via MiSeq (Illumina). Libraries were diluted to 2 nM individually then pooled for denaturation and further dilution to a final concentration of 10 pM. A MiSeq Reagent Nano Kit v2 (500-cycles) was then used for setting up the sequencing run (2x 250 bp, paired-end; up to 2x 1 million reads). A variant of the SPAdes genome assembly pipeline (St. Petersburg State University, Center for Algorithmic Biotechnology) was used for assembling of the fastQ raw sequencing data to contigs with optimized settings. WGS mutation analyses were performed at Cidara using Sequencher 5.4.6 (Gene Codes Corp.).

NGS deep sequencing of passage 10 total population (TP) samples were prepared in the same manner as those used in WGS (DNA samples were generated at Cidara and provided to BATJ for sequencing). Library construction and quantification was similar as well to the WGS protocol except that ∼1,000 ng viral cDNAs per sample were used in fragmentation. The MiSeq sequencing run followed the WGS protocol with the exception that a MiSeq Reagent Kit v2 (500-cycles) was used for sequencing run (2x 250 bp, paired-end, generating up to 2x 10 million reads). Alignment analysis utilized a variant of the breseq sequence alignment pipeline (Barrick Lab) for identification of mutations relative to the reference sequences. For metagenomic samples that contained a mixed population, the Polymorphism Mode (≥1% frequency threshold) was used to predict polymorphic (mixed) mutations. Final analyses (performed by Cidara) utilized GenBank sequences for alignment that extended beyond the reading frame in both directions; thus, clipping extraneous sequences and renumbering mutation positions was required.

### Animal Experiments

All animal use was approved under ACUP EB17-033-002 (Cidara/Explora Biolabs, San Diego).

### In vivo efficacy

Mouse efficacy studies utilized female BALB/c or BALB/c SCID (Jackson Laboratories, strain # 000651 or 001803) mice (6-8 weeks of age; *n* = 5/group for all studies). Animals were anesthetized with ketamine/xylazine (100/10 mg/kg, IP) and challenged intranasally with 3x the LD_95_ of influenza A or B suspended in 30 µL of PBS. A single IM dose of CD388 was administered via the flank at a dose volume of 10 mL/kg either 2 h after or 7 days prior to viral challenge. The general health of animals was monitored, and BW recorded daily for up to 21 days after viral challenge. Moribund, or animals exhibiting 20% or more of BW loss, were recorded as a mortality. Statistical significance was determined from survival graphs relative to vehicle using the Log-rank (Mantel-Cox) test in GraphPad Prism version 6.0.

### Lung burden and cytokine analyses

Female BALB/c mice (6-8 weeks old) were challenged intranasally with 3 × 10^2^ PFU of mouse-adapted influenza A/Puerto Rico/8/1934 (H1N1). At 2 h post-challenge, mice were administered either CD388 as a single intramuscular dose (0.01 – 3 mg/kg) or oral OST at the human equivalent dose of 5 mg/kg or 10x human equivalent dose (50 mg/kg) BID x 4 days. At 4 days post-infection, mice were sacrificed by CO_2_ and both lung lobes were harvested. Lungs were homogenized with 1 mm silica beads in 1 mL PBS using a MagNA Lyser (Roche). Homogenization was carried out at 6,000 rpm for 60 s and chilled on ice for 5 min in-between runs. After lung homogenization tubes were centrifuged for 10 min at 600 × *g* and supernatant was transferred into a new tube. For PFU determination, supernatants of lung homogenate were diluted in infection buffer ranging from 10^−1^ to 10^−6^. PFU were calculated relative to the weight of the harvested lung (PFU/g lung). For cytokine analyses, supernatants of lung homogenate were serially diluted 2-fold in 96-well plates. Cytokine levels for IL-6 and MCP-1 were determined by ELISA according to manufacturer’s instructions (R&D Systems).

