## Supplementary Information for "CD388: A universally protective Drug-Fc Conjugate that targets influenza virus neuraminidase"

**Supplemental Table 1.** Influenza viruses used in this manuscript.

| Influenza virus | Subtype | Vendor | Cat# |
|---|---|---|---|
| A/WSN/1933 | H1N1 | Chanda Lab* | n/a |
| A/Puerto Rico/8/34 | H1N1 | BEI | NR-348 |
| A/Texas/36/91 | H1N1 | BEI | NR-3223 |
| A/Bayern/07/1995 | H1N1 | IRR | FR-404 |
| A/New Caledonia/20/1999 | H1N1 | IRR | FR-395 |
| A/Solomon Islands/3/2006 | H1N1 | IRR | FR-331 |
| A/Brisbane/59/2007 | H1N1 | IRR | FR-1 |
| A/Netherlands/1324/2009 | H1N1 | BEI | NR-19835 |
| A/California/04/2009 | H1N1 | BEI | NR-13658 |
| A/California/07/2009 | (H1N1)pdm09 | IRR | FR-201 |
| A/California/12/2012 | (H1N1)pdm09 | IRR | FR-1176 |
| A/Texas/23/2012 NA H275Y | H1N1pdm09 | IRR | FRS-1169 |
| A/Michigan/45/2015 | (H1N1)pdm09 | IRR | FR-1505 |
| A/Illinois/08/2018 | H1N1 | IRR | FR-1625 |
| A/Brisbane/02/2018 | (H1N1)pdm09 | CDC | n/a |
| A/Illinois/45/2019 | (H1N1)pdm09 | IRR | FRS-1748 |
| A/Alabama/03/2020 NA H275Y | (H1N1)pdm09 | IRR | FRS-1749 |
| A/Hawaii/66/2019 | (H1N1)pdm09 | CDC | n/a |
| A/Hawaii/70/2019 | (H1N1)pdm09 | CDC | n/a |
| A/Hong Kong/1/1968 | H3N2 | BEI | NR-28621 |
| A/Aichi/2/68 | H3N2 | ATCC | VR-1680 |
| A/Victoria/3/75 | H3N2 | ATCC | VR-822 |
| A/Philippines/2/1982 | H3N2 | BEI | NR-28649 |
| A/Wisconsin/67/2005 | H3N2 | IRR | FR-397 |
| A/Bethesda/956/2006 NA R292K | H3N2 | IRR | FR-1443 |
| A/Washington/01/2007 | H3N2 | IRR | FR-1176 |
| A/Texas/12/2007 NA E119V | H3N2 | IRR | FRS-1171 |
| A/Perth/16/2009 | H3N2 | IRR | FR-370 |
| A/Victoria/361/2011 | H3N2 | IRR | FR-1061 |
| A/Texas/50/2012 | H3N2 | IRR | FR-1210 |
| A/Switzerland/9715293/2013 | H3N2 | IRR | FR-1368 |
| A/Hong Kong/4801/2014 | H3N2 | IRR | FR-1453 |
| A/Pennsylvania/46/2015 | H3N2 | IRR | FRS-1752 |
| A/Washington/33/2014 NA E119V | H3N2 | IRR | FRS-1753 |
| A/Texas/71/2017 | H3N2 | IRR | FR-1622 |
| A/Singapore/INFIMH-16-0019/2016 | H3N2 | IRR | FR-1590 |
| A/Louisiana/50/2017 | H3N2 | IRR | FR-1627 |
| A/Wisconsin/04/2018 | H3N2 | IRR | FR-1178 |
| A/South Australia/34/2019 | H3N2 | CDC | n/a |

| | | | |
|---|---|---|---|
| A/Hong Kong/2671/2019 | H3N2 | CDC | n/a |
| B/Lee/1940 | B | BEI | NR-3178 |
| B/Memphis/20/1996 | B (Yamagata) | IRR | FR-486 |
| B/Memphis/20/1996 NA R152K | B (Yamagata) | IRR | FRS-1175 |
| B/Rochester/02/2001 | B (Yamagata) | IRR | FR-1172 |
| B/Rochester/02/2001 NA D198N | B (Yamagata) | IRR | FRS-1173 |
| B/Malaysia/2506/2004 | B (Victoria) | Chanda Lab* | n/a |
| B/Florida/4/2006 | B (Yamagata) | BEI | NR-41795 |
| B/Brisbane/60/2008 | B (Victoria) | BEI | NR-42005 |
| B/Wisconsin/01/2010 | B (Yamagata) | IRR | FR-806 |
| B/Massachusetts/2/2012 | B (Yamagata) | IRR | FR-1196 |
| B/Phuket/3073/2013 | B (Yamagata) | IRR | FR-1364 |
| B/Laos/0080/2016 | B (Victoria) | IRR | FRS-1756 |
| B/Laos/0654/2016 NA H134N | B (Victoria) | IRR | FRS-1757 |
| B/Colorado/6/2017 | B (Victoria) | IRR | FR-1588 |
| B/North Carolina/25/2018 | B (Victoria) | IRR | FRS-1746 |
| B/Missouri/12/2018 NA D197E | B (Victoria) | IRR | FRS-1747 |
| B/Washington/02/2019 | B (Victoria) | CDC | n/a |
| A/Vietnam/1194/2004 | H5N1 | ViroClinics | n/a |
| A/Indonesia/05/2005 | H5N1 | ViroClinics | n/a |
| A/turkey/Turkey/1/2005 | H5N1 | ViroClinics | n/a |
| A/Hong Kong/156/97 | H5N1 | ViroClinics | n/a |
| A/Anhui/1/2013 | H7N9 | ViroClinics | n/a |

**Supplemental Table 2.** Recombinant NA cell lysates used in this manuscript.

| Recombinant Neuraminidase | Subtype | Vendor | Cat# |
|---|---|---|---|
| A/Anhui/1/2005 | H5N1 | Sino Biological | 11676-VNAHC |
| A/Shanghai/1/2013 | H7N9 | Sino Biological | 40109-VNAHC |
| A/Anhui/1/2013 | H7N9 | Sino Biological | 40108-VNAHC |

**SUPPLEMENTARY METHODS**

**Synthesis of novel ZAN dimer**

The route for synthesis of the novel ZAN dimer is described in detail below.

**Synthesis of intermediate 6**

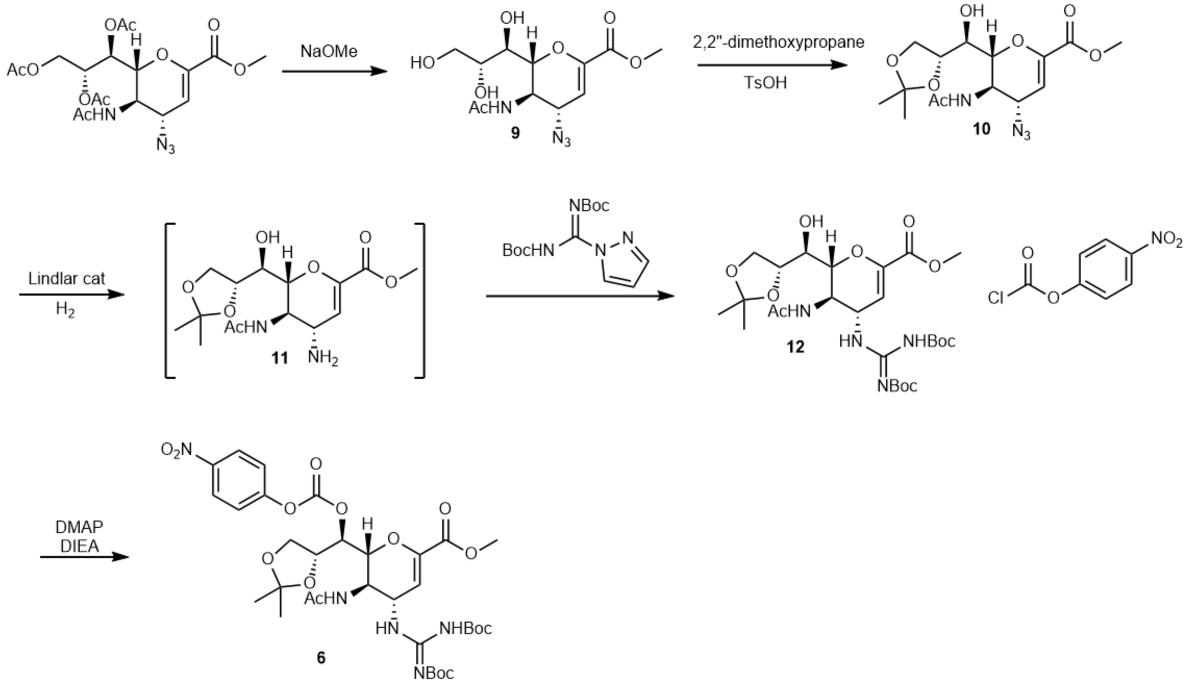


**Synthesis of 9**

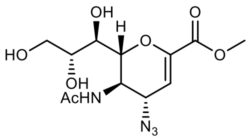

Methyl 5-acetamido-7,8,9-tri-O-acetyl-2,6-anhydro-4-azido-3,4,5-trideoxy-D-glycero-D-galacto-non-2-enonate (10.0 g, 22 mmol) was dissolved in 60 mL dry methanol, then treated with sodium methoxide in methanol (8.8 mL of 0.5 M in methanol, 4.4 mmol) while cooling in an ice-water bath. Progress of the reaction was monitored by LCMS, which was complete after 2 h. The pH of the reaction solution was then adjusted to a value of 5 to 6 by using Amberlite IRN-77 ion exchange resin. The mixture was filtered to remove the resin and evaporated to dryness under vacuum. The resulting oil was used in the next step without further purification. $^{1}$H-NMR (300 MHz, MeOD): δ 5.93 (d, *J* = 2.6 Hz, 1H), 4.39-4.09 (m, 3H), 3.92-3.82 (m, 2H), 3.80 (s, 3H), 3.71-3.58 (m, 2H), 2.04 (s, 3H); $^{13}$C-NMR (75 MHz, MeOD): δ 174.4, 163.7, 146.8, 108.4, 78.1, 71.1, 69.6, 64.8, 59.8, 53.0. 49.5, 22.7; HRMS (*m/z*): $[M+H]^+$ calcd for $C_{12}H_{19}N_4O_7$, 331.1248; found, 331.1251.

**Synthesis of 10**

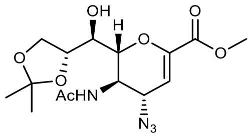


To a solution of crude **9** (0.22 mmol) in acetone (70 mL) were added 2,2-dimethoxypropane (30 mL) and p-toluenesulfonic acid monohydrate (400 mg, 2.0 mmol). The resulting solution was stirred at room temperature overnight, at which time, sodium bicarbonate (170 mg, 2.0 mmol) was added, and the mixture was concentrated to dryness. The resulting material was used in next step without further purification. $^{1}$H-NMR (300 MHz, MeOD): δ 5.93 (d, *J* = 2.5 Hz, 1H), 4.39-4.26 (m, 2H), 4.22-3.99 (m, 4H), 3.80 (s, 1H), 3.62 (dd, *J* = 1.0 Hz, 7.8 Hz, 1H), 3.35 (s, 3H), 2.03 (s, 3H), 1.37 (s, 3H), 1.33 (s, 3H); $^{13}$C-NMR (75 MHz, MeOD): δ 174.2, 163.5, 146.7, 110.3, 108.5, 78.4, 76.1, 70.5, 67.9, 59.6, 53.0, 49.5, 27.2, 25.6, 22.7; HRMS (*m/z*): $[M+H]^+$ calcd for $C_{15}H_{23}N_4O_7$, 371.1561; found, 371.1557

**Synthesis of 11 and 12**

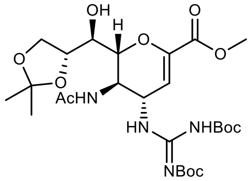


To a solution of crude **10** (0.22 mmol) in methanol (60 mL) was added Lindlar catalyst (5.0 g). The resulting mixture was vacuum flushed with hydrogen every 30 minutes to remove nitrogen gas and stirred for a total 5 h. After complete reaction as determined by LCMS, the catalyst was removed by filtration through celite. The filtrate was concentrated and used in the next step without further purification. $^{1}$H-NMR (300 MHz, MeOD): δ 5.95 (d, *J* = 2.5 Hz, 1H), 4.36-4.28 (m, 1H), 4.18-4.11 (m, 1H), 4.04-3.97 (m, 2H), 3.88 (t, *J* = 9.0 Hz, 1H), 3.78 (s, 3H), 3.62-3.56 (m, 2H), 2.06 (s, 3H), 1.37 (s, 3H), 1.33 (s, 3H); $^{13}$C-NMR (75 MHz, MeOD): δ 175.0, 164.2, 145.0, 113.9,

110.3, 78.6, 76.1, 71.2, 68.0, 52.7, 52.1, 50.4, 27.2, 25.6, 22.7; HRMS (*m/z*): $[M+H]^+$ calcd for $C_{15}H_{25}N_2O_7$, 345.1656; found, 345.1653

Crude **11** (0.22 mmol) in THF (60 mL) was treated with N,N'-Di-Boc-1H-pyrazole-1-carboxamidine (9.30 g, 30.0 mmol), and DIEA (9.9 mL, 57.0 mmol). The resulting solution was stirred at room temperature until guanylation was complete as determined by LCMS (4 h). The solution was concentrated and purified by flash chromatography eluting with 20% to 80% ethyl acetate/dichloromethane. Yield 8.8 g, 59 % for four steps. $^1$H-NMR (300 MHz, $CDCl_3$): δ 11.32 (s, 1H) 8.63 (d, $J$ = 7.7 Hz, 1H), 8.04 (s, 1H), 5.74 (d, $J$ = 1.9 Hz, 1H), 5.23-5.09 (m, 2H), 4.37-4.28 (m, 1H), 4.14-3.85 (m, 4H), 3.72 (s, 3H), 3.47-3.41 (m, 1H), 1.95 (s, 3H), 1.45 (s, 9H), 1.43 (s, 9H), 1.36 (s, 3H), 1.30 (s, 3H); $^{13}$C-NMR (75 MHz, $CDCl_3$): δ 174.0, 161.9, 161.8, 157.4, 152.6, 146.8, 109.1, 106.7, 84.5, 80.4, 78.4, 74.0, 69.7, 67.4, 53.4, 51.8, 48.5, 28.2, 28.0, 27.1, 25.2, 22.9; HRMS (*m/z*): $[M+H]^+$ calcd for $C_{26}H_{43}N_4O_{11}$, 587.2923; found, 587.2917

**Synthesis of 6**

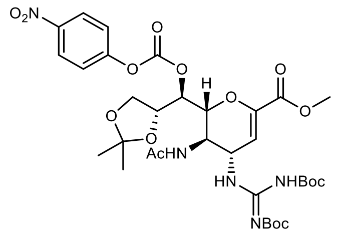


A reaction flask containing **12** (24.7 g, 42.14 mmol) dissolved in anhydrous dichloromethane (500 mL), was vacuum flushed with nitrogen. The solution was cooled in an ice-water bath, then treated with DIEA (22.1 mL, 126.42 mmol), DMAP (5.15 g, 42.14 mmol), followed by 4-nitrophenylchloroformate (17.0 g, 84.27 mmol) in portions. The solution was stirred at 0°C for 1 h then allowed to warm to room temperature for 12 h. The reaction was concentrated and purified by flash chromatography eluting with 5% to 50% ethyl acetate/dichloromethane. Yield 19.0 g, 60%. $^1$H-NMR (300 MHz, $CDCl_3$): δ 11.32 (s, 1H), 8.50 (d, $J$ = 6.8 Hz, 1H), 8.21 (d, $J$ = 9.4 Hz, 2H), 7.49 (d, $J$ = 9.4 Hz, 2H), 6.48 (d, $J$ = 8.9 Hz, 1H), 5.86 (d, $J$ = 2.1 Hz, 1H), 5.25-5.20 (m, 1H), 5.18-5.09 (m, 1H), 4.44-4.31 (m, 2H), 4.24-4.06 (m, 3H), 3.76 (s, 3H), 1.87 (s, 3H), 1.45-1.41 (m, 18H), 1.36 (s, 3H), 1.32 (s, 3H); $^{13}$C-NMR (75 MHz, $CDCl_3$): δ 171.5, 162.7, 161.6, 157.2, 155.8, 152.7, 152.5, 145.4, 145.3, 125.2, 122.4, 109.2, 108.9, 84.0, 79.8, 77.5, 75.1, 74.2,

65.6, 52.5, 48.6, 48.5, 28.2, 28.0, 26.4, 25.5, 23.1; HRMS (*m/z*): $[M+H]^+$ calcd for $C_{33}H_{46}N_5O_{15}$, 752.2985; found, 752.2983.

**Synthesis of 8**

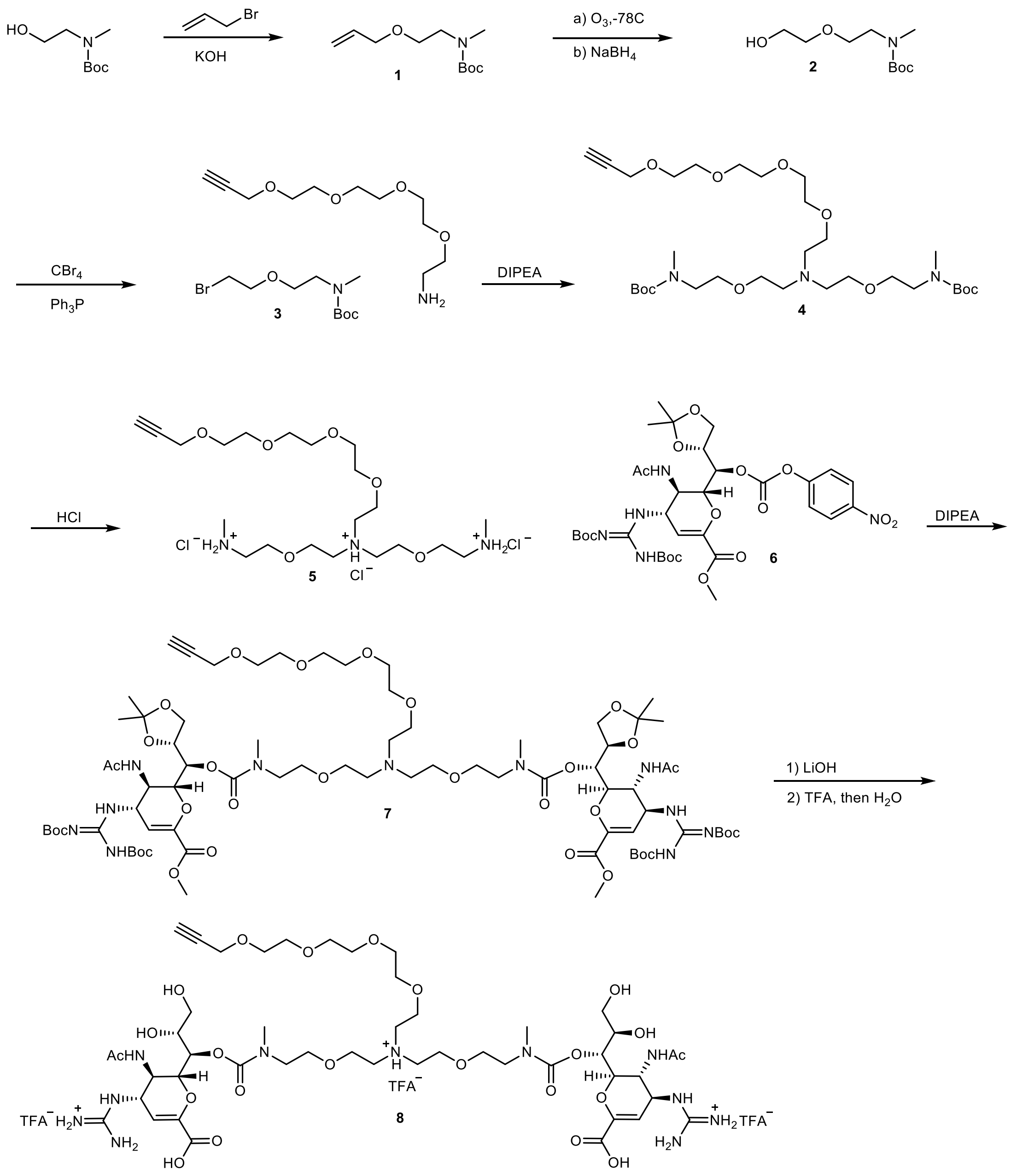

**Synthesis of 1**

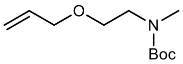


To a well-stirred solution of N-Boc-N-Me-glycinol (3.5 g, 20 mmol) in DMSO (20 mL) cooled with an ice-water bath was added allyl bromide (3.6 g, 30.0 mmol), followed by finely ground KOH powder (3.5 g, 30.0 mmol) over 15 min. The resulting solution was stirred overnight at room temperature. The resulting mixture was partitioned between 5% aq. HOAc (50 mL) and ethyl acetate (200 mL). The organic layer was separated, washed with brine, dried with sodium sulfate, filtered, and concentrated, then purified by flash chromatography eluting with 10% to 80% ethyl acetate/hexane. Yield of product 4.1g, 95%. $^{1}$H-NMR (300 MHz, MeOD): δ 5.96-5.83 (m, 1H), 5.29 (dq, $J$ = 17.2, 1.8 Hz, 1H), 5.17 (d, $J$ = 10.4 Hz, 1H), 3.98 (d, $J$ = 5.9 Hz, 2H), 3.56 (t, $J$ = 5.6 Hz, 2H), 3.40 (t, $J$ = 5.5 Hz, 2H), 2.90 (s, 3H), 1.47 (s, 9H). $^{13}$C-NMR (75 MHz, MeOD): δ 157.7, 136.2, 117.0, 81.1, 72.9, 69.4, 36.1, 35.7, 28.9. HRMS (*m/z*): $[M+Na]^+$ calculated for $C_{11}H_{21}NO_3$, 238.1419; found, 238.1415.

**Synthesis of 2**

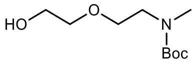


Ozone was bubbled through a solution of **1** (8.0 g, 37 mmol) in MeOH (50 mL) and DCM (50 mL) at -78 °C until the appearance of a light blue color. Unreacted ozone was removed by bubbling with oxygen for 10 min. before the addition of $NaBH_4$ (1.6 g, 40 mmol) in small portions over 10 min. After all $NaBH_4$ was added, the mixture was gradually warmed to room temperature. The resulting solution was partitioned between ethyl acetate (100 mL) and brine (50 mL). The organic layer was separated, washed with brine, dried with sodium sulfate, filtered, concentrated to an oil, and then purified by flash chromatography eluting with 10% to 80% ethyl acetate/dichloromethane. Yield of product 5.0 g, 62%. $^{1}$H-NMR (300 MHz, MeOD): δ 3.70 (t, $J$ = 4.7 Hz, 2H), 3.60-3.53 (m, 4H), 3.39 (t, $J$ = 5.6 Hz, 2H), 2.87 (s, 3H), 2.47 (bs, 1H), 1.44 (s, 9H). $^{13}$C-NMR (75 MHz, MeOD): δ 156.2, 155.9, 79.7, 72.4, 69.5, 69.1, 61.8, 48.7, 48.2, 35.3, 28.5. HRMS (*m/z*): $[M+H]^+$ calcd for $C_{10}H_{21}NO_4$, 220.1549; found, 220.1543.

**Synthesis of 3**

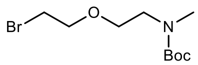


To a solution of **2** (4.4 g, 20 mmol) and $CBr_4$ (10.0 g, 30.0 mmol) in DCM (50 mL) cooled in an ice bath, was added $PPh_3$ (8.0 g, 30 mmol) slowly over 15 min. (exothermic). During the course of the addition, the internal temperature was kept below 30°C. After addition of $PPh_3$ the reaction was stirred overnight at room temperature. The crude reaction was concentrated to an oil, then purified by normal phase chromatography, eluting with 10% ethyl acetate/hexanes to 80% ethyl acetate/hexanes. Fractions containing oil droplets on the inside of the collection tubes were checked by LCMS, then combined and concentrated to a colorless oil. 4.0 g, 70.5%. $^{1}$H-NMR (300 MHz, MeOD): δ 3.76 (t, $J$ = 6.1 Hz, 2H), 3.60 (bs, 2H), 3.44 (t, $J$ = 6.1 Hz, 2H), 3.90 (bs, 2H), 2.92 (s, 3H), 1.45 (s, 9H). $^{13}$C-NMR (75 MHz, MeOD): δ 155.9, 79.5, 70.9, 69.7, 48.8, 48.4, 35.9, 35.5, 30.6, 28.6. HRMS (*m/z*): $[M+H]^+$ calcd for $C_{10}H_{20}BrNO_3$, 282.0750; found, 282.0699.

**Synthesis of 4**

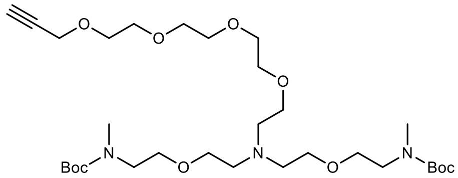


A solution of **3** (4.4 g, 15.5 mmol), propargyl-PEG4-amine (1.5 g, 6.4 mmol), and DIPEA (3.3 g, 25.8 mmol) in DMF (20 mL) were heated in an oil bath at 75°C for 18 h. The mixture was filtered, concentrated, and purified by RPLC (5% ACN/water to 100% ACN). Yield 3.85 g, 92 %. $^{1}$H-NMR (300 MHz, CDCl3): δ 4.17 (d, $J$ = 2.5 Hz, 2H), 3.87-3.79 (m, 6H), 3.67-3.61 (m, 10H), 4.59 (d, $J$ = 3.7 Hz, 2H), 3.55-3.43 (m, 10H), 3.35 (bs, 4H), 2.84 (s, 6H), 2.44 (t, $J$ = 2.4 Hz, 1H), 1.42 (s, 18H). $^{13}$C-NMR (75 MHz, CDCl3): δ 156.0, 79.8, 79.7, 74.9, 70.7, 70.6, 70.5, 70.4, 70.3, 69.2, 69.1, 66.7, 66.5, 65.9, 65.7, 65.5, 58.5, 54.5, 54.1, 48.6, 48.0, 35.3, 28.6. HRMS (*m/z*): $[M+H]^+$ calcd for $C_{31}H_{59}N_3O_{10}$, 634.4279; found, 634.4269.

**Synthesis of 5**

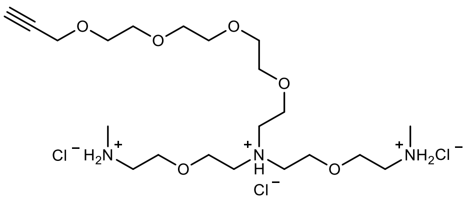


Intermediate **4** (3.85, 6.1 mmol) was treated with HCl (4 N in dioxane, 15 mL) for 2 h at room temperature. The solvent was removed by rotary evaporation, and the remaining oil was dissolved in DI water (20 mL) frozen, and lyophilized to afford the product as a light yellow oil. Yield was quantitative. $^{1}$H-NMR (300 MHz, MeOD): δ 4.21 (d, *J* = 2.4 Hz, 2H), 3.97-3.94 (m, 6H), 3.83 (t, *J* = 4.7 Hz, 4H), 3.69-3.56 (m, 18H), 3.29-3.25 (m, 4H), 2.92 (t, *J* = 2.4 Hz, 1H), 2.76 (s, 6H). $^{13}$C-NMR (75 MHz, MeOD): δ 80.8, 76.4, 71.7, 71.6, 71.5, 71.4, 70.3, 67.0, 65.9, 59.2, 55.1, 54.3, 49.8, 33.8. HRMS (*m/z*): $[M+H]^+$ calcd for $C_{21}H_{43}N_3O_6$, 434.3230; found, 434.3218.

**Synthesis of 7**

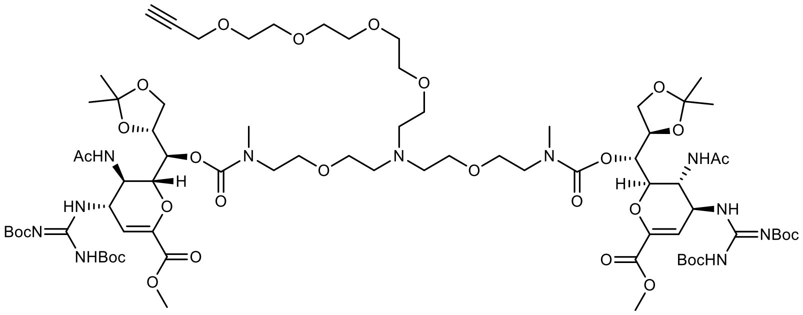


To a solution of **5** (0.68 g, 1.34 mmol) and DIPEA (0.87 g, 6.7 mmol) dissolved in anhydrous DMF (5 mL) was added **6** (2.1 g, 2.8 mmol) in portions over 1 h. The reaction was stirred at room temperature overnight, then concentrated and purified by flash chromatography, eluting with 0% to 10% methanol/dichloromethane. Yield 1.45 g, 67%. $^{1}$H-NMR (300 MHz, MeOD): δ 5.98-595 (m, 2H), 5.34 (m, 2H), 4.98-4.95 (m, 2H), 4.44 (q, *J* = 5.1 Hz, 2H), 4.37-4.32 (m, 2H), 4.24-4.15 (m, 4H), 4.19 (d, *J* = 2.5 Hz, 2H), 4.06-4.03 (m, 2H), 3.92-3.84 (m, 6H), 3.81 (s, 6H), 3.72-3.63 (m, 18H), 3.62-3.57 (m, 8H), 3.03-2.92 (m, 6H), 2.89 (t, *J* = 2.4 Hz, 1H), 1.91 (s, 6H), 1.52 (s, 18H), 1.47 (s, 18H), 1.36-1.31 (m, 12H). $^{13}$C-NMR (75 MHz, MeOD): δ 173.6, 173.4, 165.0, 164.2, 163.5, 158.0, 157.3, 156.7, 154.0, 146.1, 145.9, 111.9, 111.6, 110.1, 110.0, 85.0, 81.0, 79.0, 76.7, 76.6, 76.4, 72.2, 72.0, 71.7, 71.5, 70.9, 70.3, 66.9, 65.8, 59.2, 55.1, 53.2, 53.1, 51.6, 50.3,

48.1, 47.1, 37.1, 36.9, 36.1, 31.8, 28.6, 28.4, 28.3, 27.0, 25.7, 25.4, 23.2, 23.0. HRMS (*m/z*): $[M+H]^+$ calcd for $C_{75}H_{123}N_{11}O_{30}$, 1658.8515; found, 1658.8505.

**Synthesis of 8**

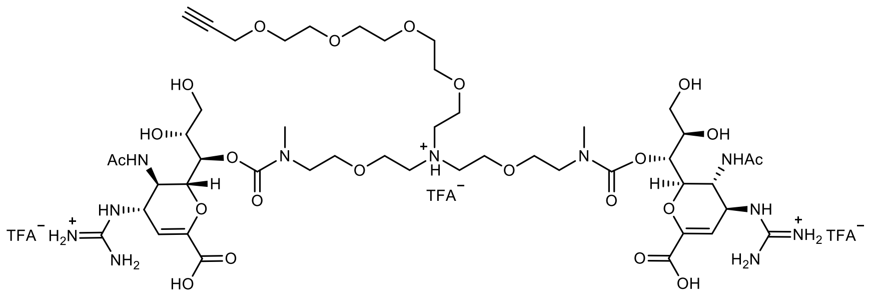


Intermediate **7** (1.45 g, 0.87 mmol) was dissolved into 3 ml MeOH then treated with a solution of lithium hydroxide (90 mg, 3.8 mmol) dissolved in deionized water (6 mL). The reaction was stirred for 15 min. at room temperature at which time LCMS showed the reaction was complete. The pH of the reaction solution was adjusted to a value of 5 to 6 by using Amberlite IRN-77 ion exchange resin, then filtered to remove the resin. The filtrate was concentrated to dryness by rotary evaporation and used in the next step without further purification. Ion found by LCMS: $[(M + 2H)/2]^+ = 815.9$.

The hydrolysis product was dissolved in dichloromethane (5 mL) and TFA (10 mL) and stirred at room temperature. The progress of the reaction was monitored by LCMS. After complete Boc-removal (~4 h), the solution was concentrated to dryness with a rotary evaporator, and then dissolved in 8 mL water. The resulting solution was stirred for another 2 hr at room temperature at which time LCMS showed complete removal of the acetonide protecting groups. This mixture was concentrated and purified by reverse phase liquid chromatography (RPLC) using an Isco CombiFlash liquid chromatograph eluted with 5% to 40% acetonitrile/water with 0.1% TFA as the modifier. Yield for three steps 780 mg, 65.0 %. $^1$H-NMR (300 MHz, MeOD): δ 5.94 (d, *J* = 2.5 Hz, 2H), 5.05 (dt, *J* = 8.9, 3 Hz, 2H), 4.59 (d, *J* = 9.9 Hz , 2H), 4.46 (dt, *J* = 8.9, 2.4 Hz , 2H), 4.29-4.19 (m, 2H), 4.22 (d, *J* = 2.4 Hz, 2H), 4.08-3.98 (m, 2H), 3.93-3.84 (m, 6H), 3.79-3.47 (m, 28H), 3.30-3.21 (m, 2H), 3.02-2.93 (m, 6H), 2.92 (t, *J* = 2.4 Hz, 1H), 1.97 (s, 6H). $^{13}$C-NMR (75 MHz, MeOD): δ 173.7, 173.6, 165.1, 157.7, 157.3, 147.3, 109.3, 80.8, 77.8, 76.3, 72.3, 71.6, 71.5, 71.4, 70.7, 70.5, 70.2, 70.0, 66.7, 65.9, 65.7, 64.6, 59.2, 59.1, 52.7, 52.6, 36.4, 35.6, 23.1, 23.0. HRMS (*m/z*): $[M+H]^+$ calcd for $C_{47}H1_{79}N_{11}O_{22}$, 1150.5479; found, 1150.5497.

**Cloning, Expression and Purification of the N-terminal extended Fc construct**. Based on EU numbering, a DNA sequence encoding residues 201 to 447 of the human IgG1 (with N-terminal extension) were used to generate the data described herein (**Sequences below**). The unpaired cysteine at position 220 in human IgG1 was mutated to serine and a M252Y/S254T/T256E (YTE) triple mutation (highlighted in bold) to alleviate issues with aggregation of the final product and extend the circulating half-life in humans, respectively. The CH1 domain residues included in the constructs are underlined. The genes for both sequences, which also included an N-terminal mouse IgG VH region secretion signal sequence for expression in Chinese hamster ovary (CHO, Gibco) cells, were cloned into the pcDNA3.1(+) vector (GenScript). The constructed plasmid was transfected into suspension ExpiCHO cells (Gibco) per manufacturer's protocol and harvested after 13 days. The Fc protein was purified using protein A affinity chromatography (MabSelect Prism A, Cytiva) and dialyzed into PBS pH 7.4.

Sequence:

NVNHKPSNTKVDKKVEPKS**S**DKTHTCPPCPAPELLGGPSVFLFPPKPKDTL**Y**I**T**R**E**PEVCVVVD

VSHEDPEVKFNWYVDGVEVHNAKTKPREEQYNSTYRVVSVLTVLHQDWLNGKEYKCKVSNKALP

APIEKTISKAKGQPREPQVYTLPPSRDELTKNQVSLTCLVKGFYPSDAVEWESNGQPENNYKTT

PPVLDSDGSFFLYSKLTVDKSRWQQGNVFSCSVMHEALHNHYTQKSLSLSPGK

**Purification of CD388**. CD388 was purified via protein A affinity chromatography and dialyzed into 100 mM EDTA in PBS pH 7.4 to remove copper and then dialyzed into PBS pH 7.4 to remove EDTA. The conjugate was purified using size exclusion chromatography (SEC) (HiLoad 26/600 Superdex, Cytiva) in PBS pH 7.4.

**Synthesis of azido-Peg Fc**. Azido-PEG4-NHS ester (0.361 mL of 0.050 M solution in DMF, 0.0181 mmol, 16 equivalents) was added to a solution of the Fc domain described above (5.3 mL, 0.0657 g, 0.00113 mmol) in PBS adjusted to pH 8.5 with 0.10 M sodium bicarbonate. LCMS (ESI) analysis was performed to monitor azide incorporation (approximately 2 h) to achieve DAR values typically between 4.5 to 5.0. Acetic acid (2.0 M) was added to lower the pH to 5.5. The resulting solution was dialyzed into 20 mM MES buffer with 100 mM NaCl at pH 6.1 (3 times, total 8-million-fold dilution). The yield of azido functionalized Fc was typically >90% as determined by UV/Vis spectrophotometry.

**Synthesis of CD388**. In a separate vial, an aqueous solution of BTTAA (6.91 mL, 1.727 mmol, 125 equivalents, 0.25 M) was mixed with aqueous solutions of $CuSO_4$ (3.45 mL, 0.345 mmol, 25 equivalents, 0.100 M), $ZnCl_2$ (1.04 mL, 1.036 mmol, 75 equivalents, 1.00 M), and 2-aminoguanidine hydrochloride (3.45 mL, 3.45 mmol, 250 equivalents, 1.00 M). In a 50 mL polypropylene conical centrifuge tube charged with a MES buffer (pH 6.1) solution of azido Fc at DAR between 4.5 to 5.0 (63.0 mL, 0.800 g, 0.0138 mmol, 1.0 equivalent), was added **8** as a solid (237 mg of triple TFA salt, 0.159 mmol, 11.5 equivalents), which dissolved readily. The copper/ligand solution was added to freshly prepared sodium ascorbate solution (3.45 mL, 3.45 mmol, 1.00 M). The pH of the homogeneous reaction mixture after combining all the reagents was approximately 4.0. The reaction was gently rotated for 3 h and then quenched by adding an aqueous solution of EDTA (14 mL, 7.0 mmol, 0.5 M, pH 8.0). The reaction was dialyzed into 100 mM EDTA in PBS (pH 7.4) to remove copper and then into PBS (pH 7.4) to remove EDTA. The crude conjugate was purified by affinity chromatography over a protein A column. The desired fractions were combined and sterile filtered and then further purified by SEC (HiLoad 26/600 Superdex, Cytiva) eluting with PBS buffer (pH 7.4). MALDI-TOF analysis of the purified final conjugate showed an average mass of 64,415 Da (DAR = 4.4 (typical DAR ranges of 4.2-4.7 are reproducibly achieved using this method)); as determined by equation DAR = (mass of conjugate – mass of Fc)/(mass of TM and azido linker)). The final product was aggregate-free (99.6% monodisperse) as determined by analytical SEC. The final solution was concentrated to 24 mg/mL in PBS (pH 7.4), in a total volume of 16.7 mL. The yield was 50%. Endotoxin levels were below detectable limits (<0.5 EU/mL) and a qualitative streaking on a culture dish showed no bacterial contamination.

**Peptide mapping**. For enzymatic digestion, using both unconjugated Fc or conjugated Fc using WT hIgG1, 200 μg protein conjugate was diluted into a final volume of 15 μL with deionized water and 45 μL of 10 mM ammonium bicarbonate, pH 8, was added. After mixing, samples were incubated at 80°C for 1 min. Samples were allowed to cool down before addition of 5 μL of a 1% solution of Promega ProteaseMAX (Promega) along with 6 μL of water. Trypsin/lys-c enzyme (Promega) was reconstituted to 2 μg/μL and added 4 μL at an enzyme:substrate ratio of 1:25. Samples were mixed and incubated at 50°C for 3 h, then quenched with 10 μL of formic acid. After sample cleanup using a Waters Oasis MCX μElution SPE plate (Waters), the sorbent was conditioned with 200 μL methanol, and then equilibrated with 200 μL water. The acidified sample was loaded and washed first with 200 μL 2% formic acid in water followed by 5% methanol in water. Samples were eluted from the sorbent with 2% ammonium hydroxide in 60:40 acetonitrile:water. Samples were analyzed by HRES-LC/MS using a Waters Acquity H-Class UPLC coupled to a Waters Q-TOF Premier mass spectrometer. The UPLC conditions included

Waters Acquity CSH C18 column, 150x2.1mm, 1.7u at 40 ºC with a flow rate of 0.5 mL/min using a gradient of 0.1% formic acid in water and 0.1% formic acid in acetonitrile. The following gradient profile was used: 0-1 min 5% B, 1-50 min 5-35% B, 50-50.5 min 35-95% B, 50.5-55 min 95% B. MSe high and low energy mass spectral data were acquired over the mass range of 90-2400 Da. Data analysis was performed using Waters MassLynx and BiopharmaLynx software.

**FcRn binding**. Precision Antibody, Columbia, MD. Human FcRn at 10 µg/mL was captured on pre-conditioned biosensors. Sensors were exposed to test article at pH 5.8 or pH 7.4. The $K_{on}$ and $K_{off}$ were measured on Octet RED96 (Octet) and the dissociation constant ($K_D$) was determined by global fitting analysis based on and $K_{on}$ and $K_{off}$.

**Cytotoxicity of CD388 in human cells.** The cytotoxicity of CD388 was determined in primary human cells (stationary cells) after 24 h and in dividing cell lines after 5 days. 5 days was chosen to allow for multiple rounds of cell division and as it is the longest incubation time used in the cell-based CPE assay. To determine the $CC_{50}$ of CD388 in primary human cells, human lung fibroblast cells (HLFs) were seeded at $1 \times 10^4$ cells per well and incubated at 37ºC, 5% $CO_2$ for 18-24 h. Freshly isolated PBMCs were added at $6 \times 10^5$ cells per well. The next day, test articles (TAs) were added at concentrations ranging from 0.0001 nM (HLFs) or 1 nM (PBMCs) to 10,000 nM to a confluent monolayer (>90%). Cytotoxicity was determined after 24 h using the CellTiter-Glo® cell viability kit according to manufacturer's instructions (Promega). Luminescence was read on an EnSpire (PerkinElmer) plate reader.

To determine the $CC_{50}$ of CD388 in human cell lines, HEp-2 cells were seeded at $5 \times 10^3$ cells per well and A549 cells at $1 \times 10^4$ cells per well and incubated at 37ºC, 5% $CO_2$ for 18-24 h. The next day, TAs were added at concentrations ranging from 0.00001 nM to 10,000 nM to a confluent monolayer (<65%). Cytotoxicity was determined after 5 days using the CellTiter-Glo® kit according to manufacturer's instructions (Promega). Luminescence was read on the EnSpire (PerkinElmer) plate reader. The % cytotoxicity was calculated using the following equation: % Cytotoxicity = (1-(RLU with TA) / (RLU cells only)) × 100%. $CC_{50}$ [nM] was determined based on % cytotoxicity with non-linear regression analysis ([inhibitor] vs normalized response with variable slope) using GraphPad Prism software version 8.

### EXTENDED DATA

**Extended Data Table 1.** Universal activity of CD388 against influenza A and B in CPE ($EC_{50}$)

| Influenza strain | Subtype | CD388 $EC_{50}$ [nM] | OST $EC_{50}$ [nM] | ZAN $EC_{50}$ [nM] | BXA $EC_{50}$ [nM] |
|---|---|---|---|---|---|
| A/WSN/1933 | H1N1 | 0.01 | 29.39 | 17.22 | 1.44 |
| A/Puerto Rico/8/34 | H1N1 | 11.25 | >10000 | 1103.00 | 1.41 |
| A/Texas/36/91 | H1N1 | 1.1 | 2880.00 | 128.50 | 1.31 |
| A/Bayern/07/1995 | H1N1 | 0.47 | 1188.00 | 7482.00 | 1.66 |
| A/Solomon Islands/3/2006 | H1N1 | 0.81 | 94.35 | 185.80 | 0.20 |
| A/Netherlands/1324/2009 | H1N1 | 1.31 | 171.10 | 113.60 | 0.82 |
| A/California/04/2009 | H1N1 | 0.48 | 100.20 | 33.77 | 0.11 |
| A/California/07/2009 | (H1N1)pdm09 | 8.99 | 98.88 | 150.80 | 10.45 |
| A/California/12/2012 | (H1N1)pdm09 | 0.75 | 99.48 | 46.85 | 6.64 |
| A/Michigan/45/2015 | (H1N1)pdm09 | 3.5 | 416.70 | 594.70 | 26.55 |
| A/Illinois/08/2018 | H1N1 | 0.31 | 742.00 | 312.90 | 9.95 |
| A/Brisbane/02/2018 | (H1N1)pdm09 | 2.54 | 248.80 | 136.50 | 10.23 |
| A/Illinois/45/2019 | (H1N1)pdm09 | 0.47 | 108.30 | 72.83 | 1.72 |
| A/Hawaii/66/2019 | (H1N1)pdm09 | 0.8 | 765.80 | 128.00 | 1.10 |
| A/Hawaii/70/2019 | (H1N1)pdm09 | 0.11 | 185.30 | 100.40 | 1.28 |
| A/Hong Kong/1/1968 | H3N2 | 1.31 | 31.33 | 95.86 | 3.43 |
| A/Aichi/2/68 | H3N2 | 5.86 | 29.42 | 345.20 | 8.89 |
| A/Victoria/3/75 | H3N2 | 2.85 | 629.90 | 369.60 | 7.74 |
| A/Philippines/2/1982 | H3N2 | 0.36 | 28.24 | 65.07 | 1.89 |
| A/Washington/01/2007 | H3N2 | 0.28 | >10000 | >10000 | 1.35 |
| A/Perth/16/2009 | H3N2 | 8.53 | 1116.00 | 1814.00 | 3.21 |
| A/Victoria/361/2011 | H3N2 | 1.86 | 5204.00 | 4936.00 | 8.92 |
| A/Texas/50/2012 | H3N2 | 0.28 | 2.58 | 3.06 | 0.00 |
| A/Switzerland/9715293/2013 | H3N2 | 0.96 | >10000 | >10000 | 0.08 |
| A/Hong Kong/4801/2014 | H3N2 | 1.24 | 1215.00 | 1643.00 | 0.98 |
| A/Texas/71/2017 | H3N2 | 4.7 | >10000 | >10000 | 7.08 |
| A/Singapore/INFIMH-16-0019/2016 | H3N2 | 2.27 | 699.60 | 27.83 | 1.26 |
| A/Louisiana/50/2017 | H3N2 | 0.88 | >10000 | >10000 | 1.23 |
| A/Wisconsin/04/2018 | H3N2 | 5.27 | >10000 | >10000 | 1.05 |
| A/Hong Kong/2671/2019 | H3N2 | 1.13 | 228.50 | 2505.00 | 3.88 |
| A/South Australia/34/2019 | H3N2 | 1.16 | 7321.00 | >10000 | 0.46 |
| B/Lee/1940 | B | 0.03 | 772.60 | 1079.00 | 1.22 |
| B/Memphis/20/1996 | B (Yamagata) | 0.99 | 994.10 | 85.64 | 0.12 |

| | | | | | |
|---|---|---|---|---|---|
| B/Rochester/02/2001 | B (Yamagata) | 1.72 | 520.90 | 401.10 | 4.10 |
| B/Malaysia/2506/2004 | B (Victoria) | 0.84 | 1912.00 | 1162.00 | 29.24 |
| B/Malaysia/2506/2004 | B (Victoria) | 1.97 | 1546.00 | 329.00 | 72.74 |
| B/Florida/4/2006 | B (Yamagata) | 2.73 | 6201.00 | 110.30 | 40.10 |
| B/Brisbane/60/2008 | B (Victoria) | 5.23 | 8287.00 | 2217.00 | 7.71 |
| B/Wisconsin/01/2010 | B (Yamagata) | 1.19 | 295.00 | 14.12 | 23.06 |
| B/Massachusetts/2/2012 | B (Yamagata) | 2.93 | 4559.00 | 98.59 | 37.99 |
| B/Phuket/3073/2013 | B (Yamagata) | 0.13 | 151.90 | 93.24 | 1.28 |
| B/Laos/0080/2016 | B (Victoria) | 1.31 | 976.30 | 883.10 | 9.99 |
| B/Colorado/6/2017 | B (Victoria) | 8.71 | >10000 | 2991.00 | 2.79 |
| B/North Carolina/25/2018 | B (Victoria) | 6.76 | 6496.00 | 31.90 | 40.82 |
| B/Washington/02/2019 | B (Victoria) | 6.95 | >10000 | 954.50 | 11.44 |

**Extended Data Table 2.** Broad-spectrum activity of CD388 against high-pathogenic influenza A (H5N1) and (H7N9) in microneutralization ($EC_{50}$)

| Influenza strain | Subtype | CD388 $EC_{50}$ [nM] | OST $EC_{50}$ [nM] | ZAN $EC_{50}$ [nM] | BXA $EC_{50}$ [nM] |
|---|---|---|---|---|---|
| A/Vietnam/1194/2004 | H5N1 | 0.88 | 1.13 | 2.24 | <0.03 |
| A/Indonesia/05/2005 | H5N1 | 0.95 | 29.49 | 5.09 | <0.03 |
| A/turkey/Turkey/1/2005 | H5N1 | 0.25 | 4.06 | 0.69 | <0.03 |
| A/Hong Kong/156/97 | H5N1 | 0.09 | 4.95 | 0.92 | <0.03 |
| A/Anhui/1/2013 | H7N9 | 0.04 | 13.70 | 43.54 | <0.03 |

**Extended Data Table 3.** $CC_{50}$ of CD388 in primary human cells (PBMCs and HLFs) cells after 24 h incubation

| PMBCs donor 1 $CC_{50}$ [nM] | PMBCs donor 2 $CC_{50}$ [nM] | HLFs $CC_{50}$ [nM] |
|---|---|---|
| >10,000 | >10,000 | >10,000 |
| >10,000 | >10,000 | >10,000 |

**Extended Data Table 4.** $CC_{50}$ of CD388 in HEp-2 and A549 cells after 5 day incubation

| HEp-2 $CC_{50}$ [nM] | A549 $CC_{50}$ [nM] |
|---|---|
| >10,000 | >10,000 |

**Extended Data Table 5.** CD388 electivity index (SI) in MDCK-SIAT1 cells

| Influenza subtype | CD388 SI | OST SI | ZAN SI | BXA SI |
|---|---|---|---|---|
| A/H1N1 | >12472 | >54.0 | >77.8 | 5816.9 |
| A/H3N2 | >7846 | >8.6 | >4.6 | 5165.7 |
| B | >5804 | >6.7 | >24.9 | 646.6 |

**Extended Data Table 6.** Minimal protective dose of CD388 administered as a single IM dose in lethal BALB/c mouse models of influenza infection

| Influenza subtype | **Treatment** Minimal protective dose | **Statistical significance** | **Prophylaxis** Minimal protective dose | **Statistical significance** |
|---|---|---|---|---|
| A/WSN/1933 (H1N1) | 0.3 mg/kg | P = 0.0035 | 0.3 mg/kg | P = 0.0027 |
| A/Puerto Rico/8/1934 (H1N1) | 0.3 mg/kg | P = 0.0035 | 0.1 mg/kg | P = 0.0025 |
| A/California/07/2009 (H1N1)pdm09 | 1 mg/kg | P = 0.0027 | 1 mg/kg | P = 0.0020 |
| A/California/12/2012 (H1N1)pdm09 | 1 mg/kg | P = 0.0023 | 1 mg/kg | P = 0.0025 |
| A/North Carolina/04/2014 (H1N1)pdm09 | 1 mg/kg | P = 0.0047 | 0.1 mg/kg | P = 0.0015 |
| A/Hawaii/70/2019 (H1N1)pdm09 | 1 mg/kg | P = 0.0023 | 1 mg/kg | P = 0.0016 |
| A/Hong Kong/1/1968 (H3N2) | 0.3 mg/kg | P = 0.0135 | 0.3 mg/kg | P = 0.0002 |
| B/Florida/4/2006 (Yamagata) | 0.3 mg/kg | P = 0.0016 | 0.3 mg/kg | P = 0.0035 |
| B/Malaysia/2506/2004 (Victoria) | 0.3 mg/kg | P = 0.0031 | 1 mg/kg | P = 0.0020 |
| B/Colorado/6/2017 (Victoria) | 0.3 mg/kg | P = 0.0016 | 1 mg/kg | P = 0.0031 |

**Extended Data Table 7.** CD388 PK after IV administration in BALB/c mice or cynomolgus monkey

| Species | Dose [mg/kg] | $T_{max}$ [h] | $c_{max}$ [µg/mL] | AUC [h x µg/mL] | Calculated $T_{1/2}$ [h] |
|---|---|---|---|---|---|
| Mouse | 3 | 0.25 | 48.4 | 1,390 | 106 |
| Monkey | 10 | 0.25 | 220 | 24,600 | 364 |

**Extended Data Table 8.** Equilibrium binding $K_D$ measurement of CD388, unconjugated YTE and WT Fc to human FcRn protein at pH 5.8 and pH 7.4

| pH | CD388 $K_D$ (M) | YTE Fc $K_D$ (M) | WT Fc $K_D$ (M) |
|---|---|---|---|
| 5.8 | <1.0E-12 | <1.0E-12 | 6.92E-09 |
| 7.4 | 1.33E-07 | 2.02E-07 | 4.35E-07 |

**Extended Data Table 9.** CD388 dose-proportionality after IM administration in mice

| Test article | Dose [mg/kg] | $T_{max}$ [h] | $C_{max}$ [µg/mL] | AUC [h x µg/mL] |
|---|---|---|---|---|
| CD388 | 0.3 | 4 | 1.37 | 107 |
| | 1 | 24 | 2.77 | 270 |
| | 3 | 24 | 10.39 | 796 |
| | 10 | 4 | 24.98 | 2266 |
| | 30 | 24 | 116.09 | 6507 |

**Extended Data Table 10.** Cross-resistance and susceptibility of CD388 serial passage-selected NA mutants versus isogenic WT parents in cell-based PRA

| Test article | Drug | NA | HA | $EC_{50}$ [nM] | Fold-change |
|---|---|---|---|---|---|
| A/WSN/1933 (H1N1) | CD388 | WT | WT | 0.09 | 2.6 |
| | | S231R* | WT | 0.23 | |
| | OS | WT | WT | 80.11 | 0.6 |
| | | S231R* | WT | 50.02 | |
| | ZA | WT | WT | 6.68 | 1.5 |
| | | S231R* | WT | 9.97 | |
| A/Victoria/3/75 (H3N2) | CD388 | WT | WT | 1.65 | 2.7 |
| | | S246V | T453I | 4.49 | |
| | OS | WT | WT | 1.80 | 5.5 |
| | | S246V | T453I | 9.92 | |
| | ZA | WT | WT | 6.09 | 8.7 |
| | | S246V | T453I | 52.74 | |
| | CD388 | WT | WT | 1.65 | 2.8 |
| | | S246V | N113H/T453I | 4.66 | |
| | OS | WT | WT | 1.80 | 4.5 |
| | | S246V | N113H/T453I | 8.11 | |
| | ZA | WT | WT | 6.09 | 10.6 |
| | | S246V | N113H/T453I | 64.57 | |

*Due to a 16-AA-encoding truncation in the NA gene of A/WSN/1933, residue S231 corresponds to S247 in the N1 wild-type NA consensus sequence.

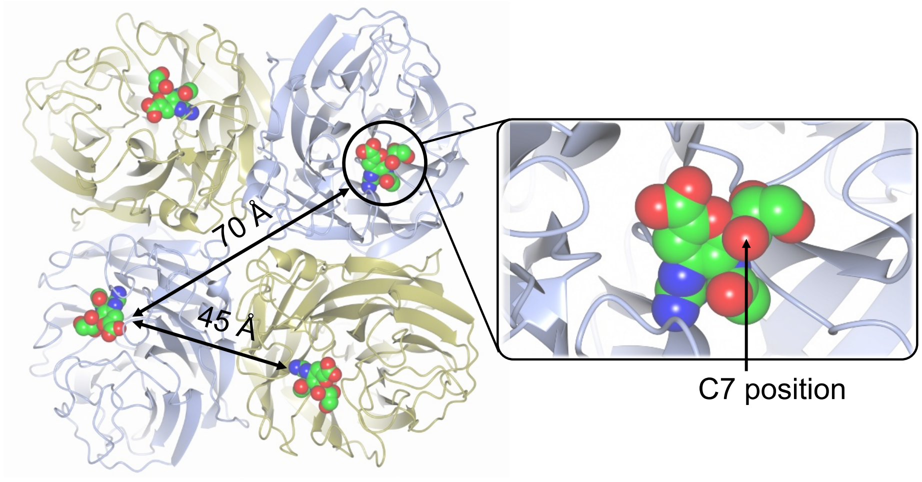


**Extended Data Figure 1. The NA tetramer with tetrameric ZAN.** Crystal structure of tetrameric ZAN (green) complexed to an NA tetramer (PDB code 3TI5) from A/California/04/2009 H1N1. The C7 position on ZAN that was chosen for attachment to the Fc

carrier (via an NHS ester) in the work presented here is highlighted. C7 on ZAN is solvent exposed and unencumbered sterically, allowing conjugation to the C7 position with minimal influence on binding affinity to NA.

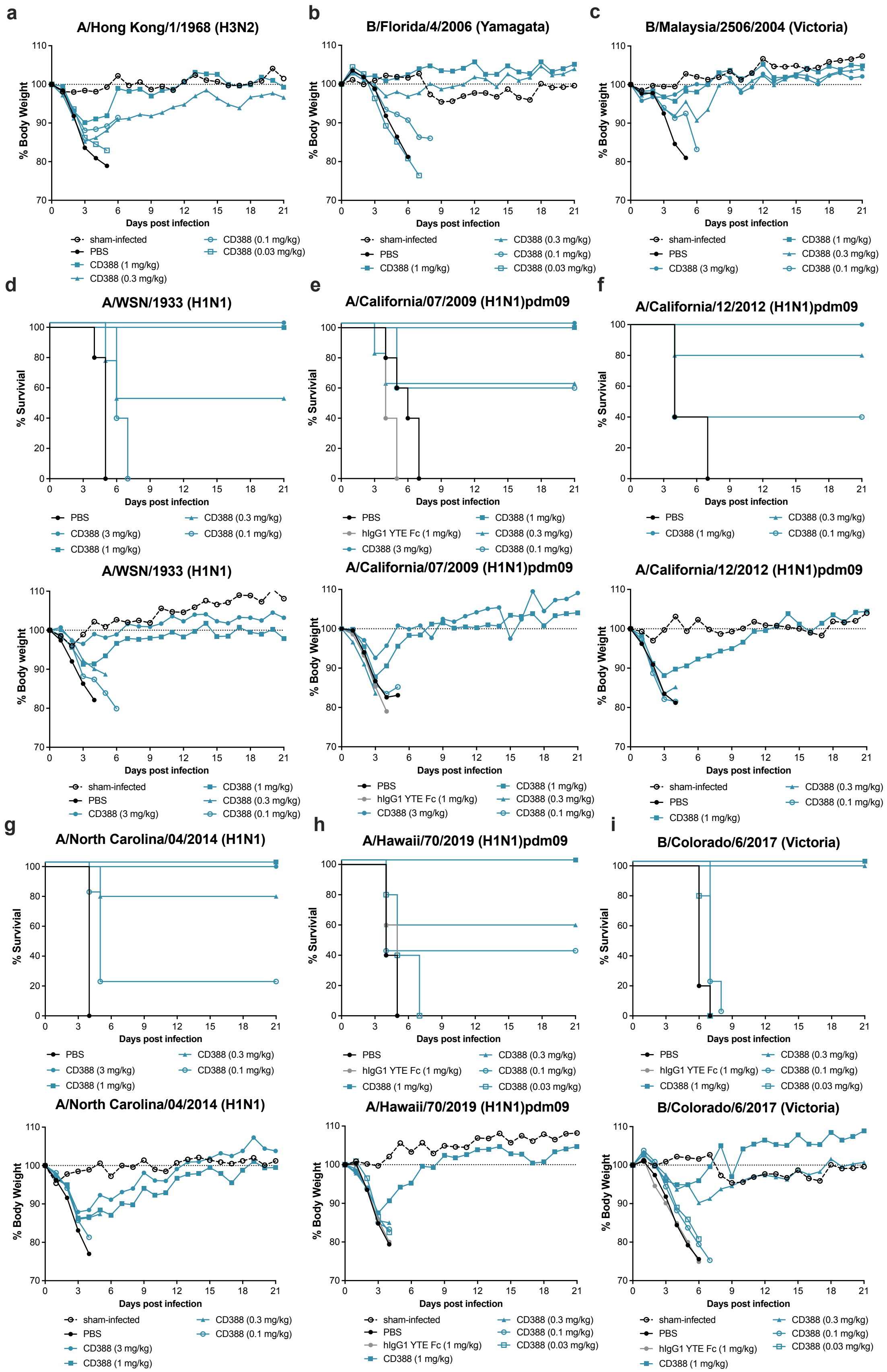

a
A/Hong Kong/1/1968 (H3N2)
% Body Weight
Days post infection
sham-infected
PBS
CD388 (1 mg/kg)
CD388 (0.3 mg/kg)
CD388 (0.1 mg/kg)
CD388 (0.03 mg/kg)
b
B/Florida/4/2006 (Yamagata)
% Body Weight
Days post infection
sham-infected
PBS
CD388 (1 mg/kg)
CD388 (0.3 mg/kg)
CD388 (0.1 mg/kg)
CD388 (0.03 mg/kg)
c
B/Malaysia/2506/2004 (Victoria)
% Body weight
Days post infection
sham-infected
PBS
CD388 (3 mg/kg)
CD388 (1 mg/kg)
CD388 (0.3 mg/kg)
CD388 (0.1 mg/kg)
d
A/WSN/1933 (H1N1)
% Survivial
Days post infection
PBS
CD388 (3 mg/kg)
CD388 (1 mg/kg)
CD388 (0.3 mg/kg)
CD388 (0.1 mg/kg)
e
A/California/07/2009 (H1N1)pdm09
% Survivial
Days post infection
PBS
hIgG1 YTE Fc (1 mg/kg)
CD388 (3 mg/kg)
CD388 (1 mg/kg)
CD388 (0.3 mg/kg)
CD388 (0.1 mg/kg)
f
A/California/12/2012 (H1N1)pdm09
% Survivial
Days post infection
PBS
CD388 (1 mg/kg)
CD388 (0.3 mg/kg)
CD388 (0.1 mg/kg)
A/WSN/1933 (H1N1)
% Body weight
Days post infection
sham-infected
PBS
CD388 (3 mg/kg)
CD388 (1 mg/kg)
CD388 (0.3 mg/kg)
CD388 (0.1 mg/kg)
A/California/07/2009 (H1N1)pdm09
% Body Weight
Days post infection
PBS
hIgG1 YTE Fc (1 mg/kg)
CD388 (3 mg/kg)
CD388 (1 mg/kg)
CD388 (0.3 mg/kg)
CD388 (0.1 mg/kg)
A/California/12/2012 (H1N1)pdm09
% Body Weight
Days post infection
sham-infected
PBS
CD388 (1 mg/kg)
CD388 (0.3 mg/kg)
CD388 (0.1 mg/kg)
g
A/North Carolina/04/2014 (H1N1)
% Survivial
Days post infection
PBS
CD388 (3 mg/kg)
CD388 (1 mg/kg)
CD388 (0.3 mg/kg)
CD388 (0.1 mg/kg)
h
A/Hawaii/70/2019 (H1N1)pdm09
% Survivial
Days post infection
PBS
hIgG1 YTE Fc (1 mg/kg)
CD388 (1 mg/kg)
CD388 (0.3 mg/kg)
CD388 (0.1 mg/kg)
CD388 (0.03 mg/kg)
i
B/Colorado/6/2017 (Victoria)
% Survivial
Days post infection
PBS
hIgG1 YTE Fc (1 mg/kg)
CD388 (1 mg/kg)
CD388 (0.3 mg/kg)
CD388 (0.1 mg/kg)
CD388 (0.03 mg/kg)
A/North Carolina/04/2014 (H1N1)
% Body Weight
Days post infection
sham-infected
PBS
CD388 (3 mg/kg)
CD388 (1 mg/kg)
CD388 (0.3 mg/kg)
CD388 (0.1 mg/kg)
A/Hawaii/70/2019 (H1N1)pdm09
% Body Weight
Days post infection
sham-infected
PBS
hIgG1 YTE Fc (1 mg/kg)
CD388 (1 mg/kg)
CD388 (0.3 mg/kg)
CD388 (0.1 mg/kg)
CD388 (0.03 mg/kg)
B/Colorado/6/2017 (Victoria)
% Body Weight
Days post infection
sham-infected
PBS
hIgG1 YTE Fc (1 mg/kg)
CD388 (1 mg/kg)
CD388 (0.3 mg/kg)
CD388 (0.1 mg/kg)
CD388 (0.03 mg/kg)

**Extended Data Figure 2. Universal activity of CD388 against influenza A and B in lethal mouse models.** CD388 efficacy against lethal challenge with influenza in BALB/c mice. Shown is dose-response % body weight change in response to CD388 ranging from 0.03 – 1 mg/kg administered IM at 2 h post-infection against lethal challenge with influenza (**a**) A/Hong Kong/1/1968 (H3N2), (**b**) B/Florida/4/2006 (Yamagata), (**c**) B/Malaysia/2506/2004 (Victoria), and % body weight change and survival against (**d**) A/WSN/1933 (H1N1), (**e**) A/California/07/2009 (H1N1)pdm, (**f**) A/California/12/2012 (H1N1)pdm09, (**g**) A/North Carolina/04/2014 (H1N1)pdm09, (**h**) A/Hawaii/70/2019 (H1N1)pdm09, and (**i**) B/Colorado/6/2017 (Victoria) in BALB/c mice.

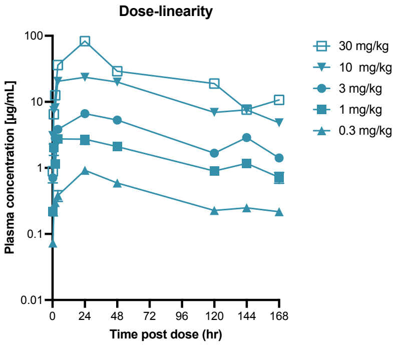


**Extended Data Figure 3. CD388 dose linearity.** CD388 plasma levels for doses of 0.3 – 30 mg/kg after IM administration in mice (mean ± SD; *n* = 2/group) as measured by Fc capture.

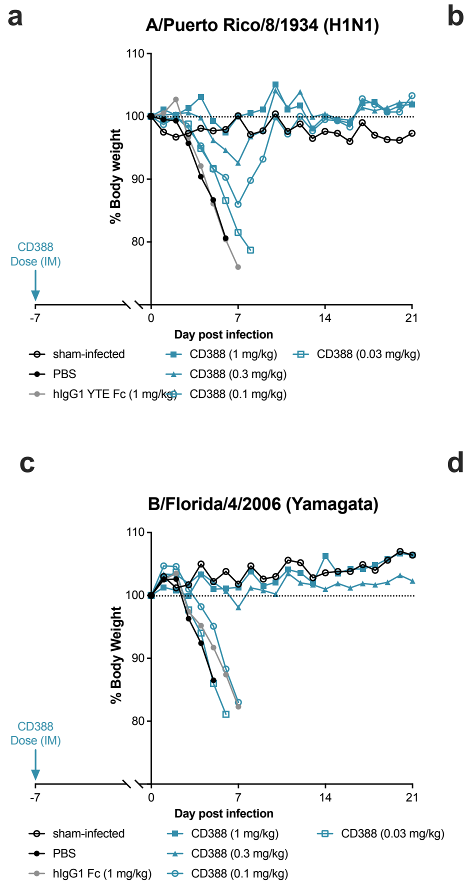


**Extended Data Figure 4. CD388 prophylactic activity – body weight change.** % body weight change in response to CD388 administered IM at 7 days prior to lethal challenge against influenza (**a**) A/Puerto Rico/8/1934 (H1N1), (**b**) A/Hong Kong/1/1968 (H3N2), (**c**) B/Florida/4/2006 (Yamagata) and (**d**) B/Malaysia/2506/2004 (Victoria).

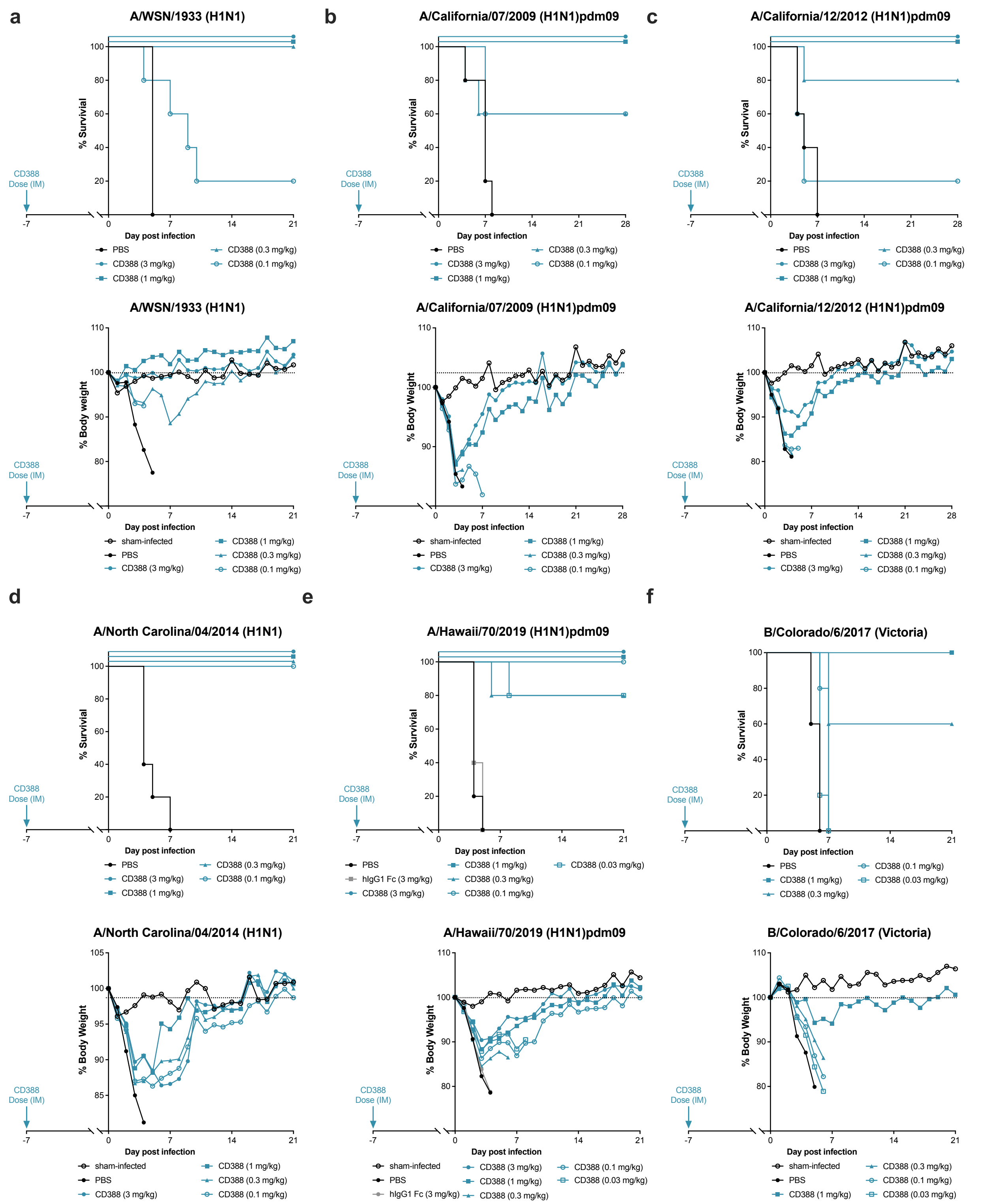


**Extended Data Figure 5. CD388 prophylactic activity – Survival and body weight change.**

Survival and % body weight change in response to CD388 administered IM at 7 days prior to lethal challenge against influenza (**a**) A/WSN/1933 (H1N1), (**b**) A/California/07/2009 (H1N1)pdm, (**c**)

A/California/12/2012 (H1N1)pdm09, (**d**) A/North Carolina/04/2014 (H1N1)pdm09, (**e**) A/Hawaii/70/2019 (H1N1)pdm09, and (**f**) B/Colorado/6/2017 (Victoria) in BALB/c mice.
